# Presence of an unfamiliar peer reverses escalated cocaine intake in male rats: involvement of the subthalamic nucleus

**DOI:** 10.1101/2023.02.07.527550

**Authors:** Cassandre Vielle, Lucie Vignal, Alix Tiran-Cappello, Julie Meffre, Nicolas Maurice, Mickael Degoulet, Cécile Brocard, Florence Pelletier, Yann Pelloux, Christelle Baunez

**Affiliations:** Institut de Neurosciences de la Timone, UMR 7289 CNRS & Aix-Marseille Université 13005 Marseille, France

**Keywords:** social context, addiction, basal ganglia, optogenetic inhibition, extended-access, social interactions

## Abstract

The immediate social context critically modulates drug consumption. The presence of an unfamiliar conspecific, naive to the drug, at the time of consumption reduces cocaine self-administration in male rats during short-access sessions, as well as drug intake in human cocaine users. The subthalamic nucleus (STN), a brain structure involved in cocaine addiction and limbic processes, has been proposed to mediate such social influence on this limited level of drug intake. Whether this influence extends to escalated drug consumption remains an open question. In this study, we compared the effect of the presence of an unfamiliar peer, naive to cocaine, on cocaine self-administration in rats having been exposed to either short (2h) or long-access sessions (6h). We showed that the presence of the peer drastically reduced both limited and escalated cocaine intake in males, while it had no effect on females during short-access sessions. Assessing the effect of STN photo-inhibition or high frequency (HF) stimulation in male rats, we demonstrated that it had no effect in the absence of the conspecific in short-access sessions, STN photo-manipulation suppressed the influence of the peer’s presence. Moreover, STN photo-inhibition and HF stimulation decreased drug consumption in long-access sessions, but no additive effect was observed when associated with the peer’s presence, confirming an overriding effect of STN manipulation. Taken together, these results highlight the potential influence of socially oriented manipulations on cocaine intake and further position the STN as a critical mediator of the effect of social presence on addictive-like behaviors.

## Introduction

Currently, no efficient pharmacological treatments exist for cocaine abuse. Behavioral and social interventions show promise in helping users control their drug consumption (1,2), emphasizing the importance of environmental factors. Despite the limited number of preclinical studies considering these influences (3–5), social factors significantly impact drug use in both rodents and humans (6,7). Positive social contexts can mitigate drug consumption (8,9), whereas adverse social interactions or isolation increase the risk of substance use disorders (10–14). Recent studies propose social contact as an alternative reward to drugs (15–19). However, most drug users consume in environments where peers are present but not offered as an alternative to drug use (for reviews, see (6,20,21)). Such proximal social factors affect drug intake in complex ways (22–27).

In male rats, as in humans, the presence of a drug-naïve unfamiliar congener reduces cocaine intake during short-access sessions (25). Although extended access to cocaine, which leads to escalated intake, can be reduced in rats reared alongside another rat without cocaine access (27), the influence of social presence at the time of consumption following escalation remains unexplored.

Understanding the interplay between social factors and neural circuitry may reveal new approaches to addiction treatment. Emerging research suggests that the Subthalamic Nucleus (STN) plays a role in the beneficial effects of social factors on limited drug use (25,26,28). Manipulating STN activity can modulate cocaine motivation (29,30) and drug-seeking behavior (31–34), making it a potential target for addiction treatment (35). The STN is also involved in social processes (25,26,28,36–38), further supporting its role in mediating the influence of social context on drug use.

Here, we first checked whether females would reduce their cocaine intake in presence of an unfamiliar congener, naïve of cocaine, the optimal characteristics for reducing cocaine intake under short-access conditions in males (25). We then tested if this social presence could reduce cocaine intake in male rats following a drug escalation procedure, which produces compulsive-like responding (39). We used a continuous reinforcement procedure with a well-established dose (26,28–31). Additionally, we examined the STN’s contribution to this social influence by employing optogenetic approaches to selectively modulate STN activity (photo-inhibition or photo-stimulation at high frequency (HF)) after escalation of drug intake.

## Materials and Methods

A detailed description of the methods is provided in the Supplementary. All animal procedures were conducted in accordance with the French regulation (Decree 2010-118) and approved by the local ethics committee and the French Ministry of Agriculture (#03129.01). Male (n=80 *i.e.* n=50 for behavioral experiment and n=30 as observer peers or for electrophysiological recordings) and female (n=13 *i.e*. n=8 for behavioral experiment and n=5 as observer peers) adult Lister Hooded rats (Charles River, Saint-Germain-sur-l’Arbresle, France) were used.

We conducted an experiment with female rats to compare their reactivity to social presence in short access condition. They underwent the schedule used formerly with males ((25); **Fig S1A**) that consisted of 2-h daily sessions of cocaine self-administration (continuous reinforcement FR1, for 250μg cocaine per 90μl infusion in each 5s injection) until consumption stabilized (<25% variability in injections for 5 consecutive days) - i.e., baseline. Then, for 5 consecutive days, females self-administered cocaine in a presence of an unfamiliar, naïve to cocaine, female during the 2-h daily sessions. Given the results (see details below), only males were used for the rest of the experimentation. Male rats underwent a stereotaxic surgery to transfect STN neurons (40) with opsin using AAV5 (UNC Vector Core, Chapel Hill, USA). Three groups of rats (EYFP-control, ARCHT3.0 and hChR2) received respectively the following constructs: CaMKII-EYFP, CaMKII-ArchT3.0-p2A-EYFP-WPRE and CaMKII-hChR2(E123T/T159C)-p2A-EYFP-WPRE. The three group (EYFP-control, ARCHT3.0, hChR2) were subjected to the procedure in parallel according to the same schedule. All male rats (**Fig.1A**) underwent the same baseline acquisition as the females. After acquiring a stable baseline level of consumption for five consecutive days, they underwent 6-h (long-access) or 2-h (short-access) daily sessions for 20 days. The short-access group served as a control for the escalation effect expected in the long-access group. Finally, the rats underwent 10 additional 2-h sessions with laser stimulation (ON), consisting of two blocks of 5 consecutive days during which rats self-administered cocaine alone or in presence of a cocaine-naïve unfamiliar male observer in the adjacent compartment of the self-administration chamber. To account for potential social recognition memory effects (38), a different observer peer was used daily. The order of *‘Without peer’ vs. ‘With peer’* conditions and the presentation of observers were counterbalanced between rats.

**Figure 1.**
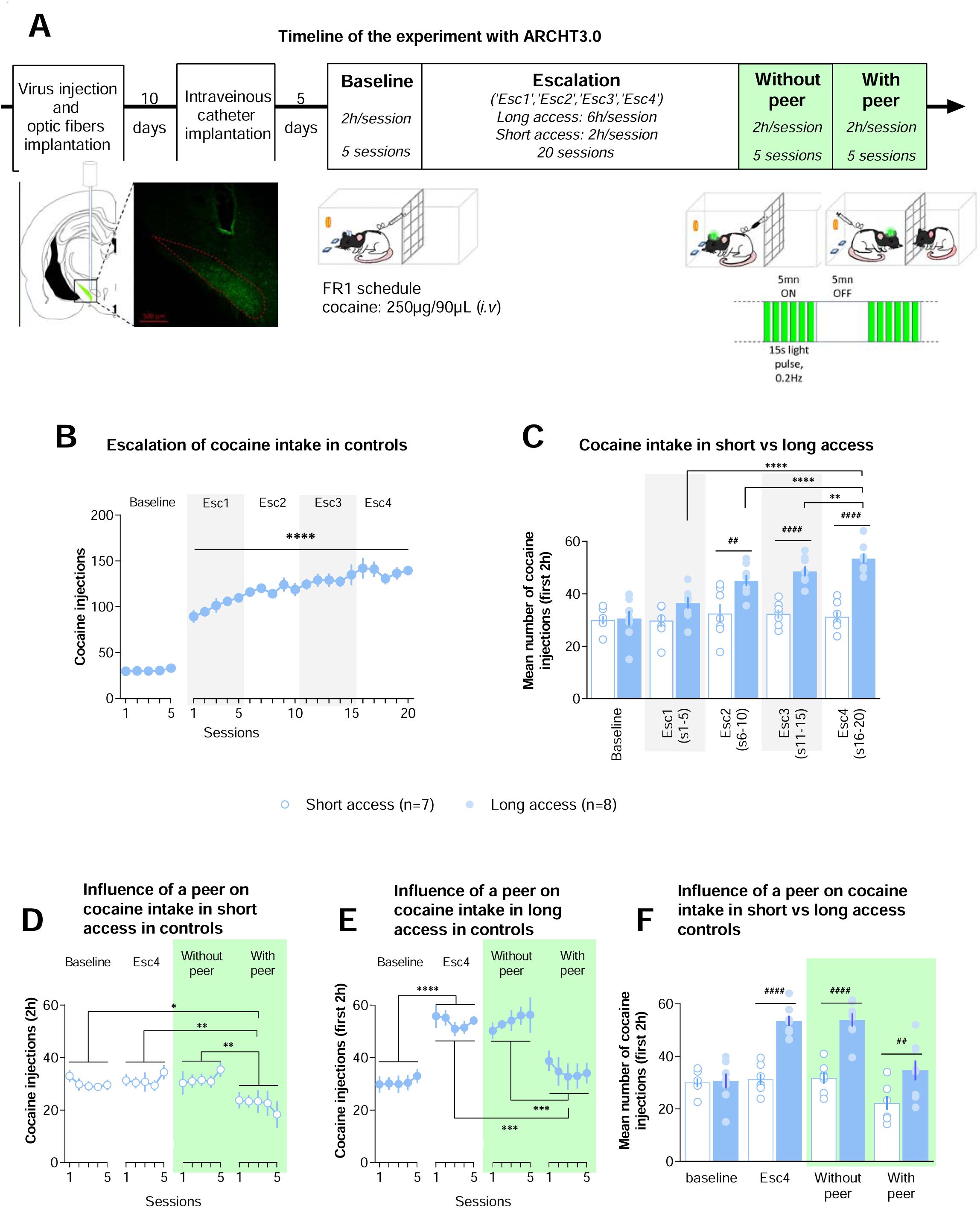
Peer’s presence reduces both limited and addiction-like cocaine intake.

The laser (532nm) period used during *‘Without peer’* and *‘With peer’* consisted in 12 blocks of 5-min ON intermingled with 12 blocks of 5 min OFF. For STN HF-stimulation (hChR2), the 5-min block ON consisted in 2ms light (10mW at the fiber tip) pulses at 130Hz, while for STN photo-inhibition (ArchT3.0) the 5-min block ON consisted of 15s light (5mW at the fiber tip) pulses at 0.2Hz.

The number of cocaine injections was analyzed by fitting linear mixed models (REstricted Maximum Likelihood (REML), to account for missing data (due to catheter disconnections) at random. Optogenetic groups (EYFP-control, ARCHT3.0, or hChR2) and access duration to cocaine (short or long-access) were considered as between factors, while sessions or conditions (baseline, escalation: ‘*Esc1*’, ‘*Esc2*’, ‘*Esc3*’, ‘*Esc4*’, *‘With peer’* and in ‘*Without peer’*) as within factors. Results are expressed as mean ±SEM. Statistical analyses details are provided in Supplementary Table 1 for behavior and Table 2 for STN electrophysiological recordings.

## Results

The results are focused only on the number of cocaine injections, since inactive lever presses did not show significant changes (REML analysis showed no significant group, session, condition effects nor interactions; **Table 1**, n=26 long access rats (8 EYFP, 9 ARCHT3.0 and 9 hChR2)).

**Table 1.**
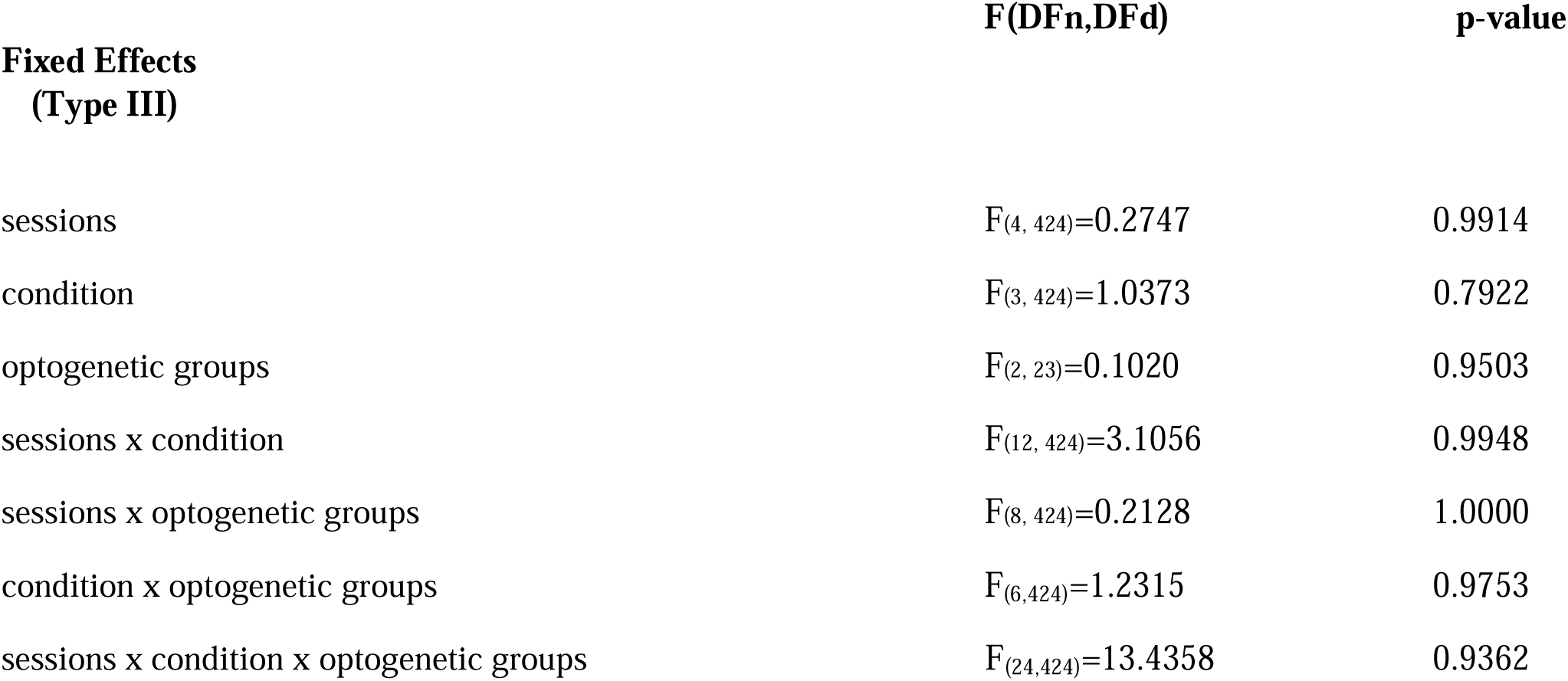
Results of the REML performed on the five sessions of baseline, *‘Esc4’*, ‘*With peer*’ and ‘*Without pee*r’ inactive lever presses in EYFP-control, ArchT3.0 and hChR2 animals subjected to the long access to the drug.

| **Fixed Effects**<br>**(Type III)** | **F(DFn,DFd)** | **p-value** |
| --- | --- | --- |
| sessions | $F_{(4, 424)}=0.2747$ | 0.9914 |
| condition | $F_{(3, 424)}=1.0373$ | 0.7922 |
| optogenetic groups | $F_{(2, 23)}=0.1020$ | 0.9503 |
| sessions x condition | $F_{(12, 424)}=3.1056$ | 0.9948 |
| sessions x optogenetic groups | $F_{(8, 424)}=0.2128$ | 1.0000 |
| condition x optogenetic groups | $F_{(6, 424)}=1.2315$ | 0.9753 |
| sessions x condition x optogenetic groups | $F_{(24, 424)}=13.4358$ | 0.9362 |

**Table 2.**
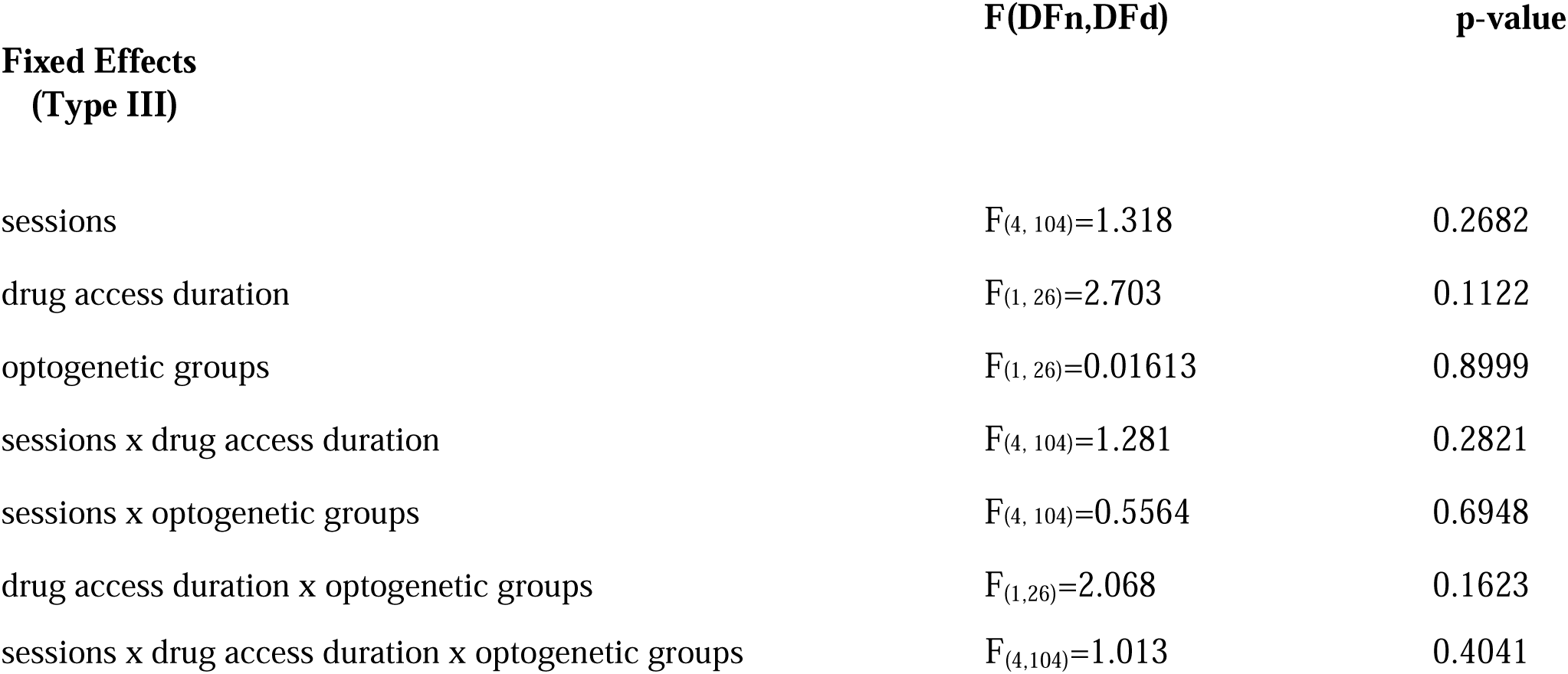
Results of the REML performed on the five sessions of baseline cocaine consumption in EYFP-control and ArchT3.0 animals subjected to short and long access to the drug.

### Female rats in short access are unsensitive to the presence of an unfamiliar, naïve to cocaine, peer

In female rats with short-access, the social presence had no effect on cocaine intake (**Fig.S1B**, REML, social condition: F(1,7)=0.0021, p=0.9634, sessions: F(4,28)=0.8872, p=0.9264, interaction: F(4,28)=1.6970, p=0.7913, Short access female control group n=8). The cocaine intake during the ‘*With peer*’ condition (38.68±1.50 injections) was similar to the one during baseline (38.55±0.39). Given this lack of effect, the rest of the experiment was conducted with male rats.

### Male rats in long-but not short-access to cocaine escalated their drug consumption

Before cocaine long-access, male rats showed a stable baseline intake (**Fig.1B**; REML, sessions: F(4,28)=1.718, p=0.174, long-access EYFP-control group n=8), consuming an average of 30.6±3.0 injections per session. Across long access sessions, their cocaine consumption significantly increased (REML, sessions: F(19,123)=6.7, p<0.0001), reaching 139.6±5.6 injections at the last session (post-hoc linear trend F(1,123)=111, p<0.0001). Animals maintained in short-access also showed a stable baseline cocaine intake (REML, sessions: F(4,24)=1.633, p=0.1984, short-access EYFP-control group n=7), consuming an average of 30.0±1.7 injections. However, their drug consumption did not increase across the following twenty short-access sessions (REML, sessions: F(19,103)=0.9675, p=0.5046) reaching 34.4±2.9 injections at the last session. To compare short and long-access intake, the cocaine injections taken in the two first hours of each session was averaged per block of 5 sessions (baseline; escalation: *‘Esc1’*: sessions 1-5; *‘Esc2’*: sessions 6-10; *‘Esc3’*: sessions 11-15 and *‘Esc4’*: sessions 16-20) and compared between rats in short-*vs.* long-access (**Fig.1C**; REML: drug access duration: F(1,13)=25.04, p=0.0002; block: F(4,52)=15.52, p<0.0001, interaction: F(4,52)=10.49, p<0.0001, long-access EYFP-control group n=8 and short-access EYFP-control group n=7). Mirroring our previous results, drug consumption during the two first hours was escalated from the baseline to *‘Esc4’* in rats subjected to long*-* access sessions (Dunnett, *‘Esc4’ vs.* baseline and *‘Esc1’* p<0.0001, ‘Esc4’ *vs. ‘Esc2’* p=0.04 and ‘Esc4’ *vs.‘Esc3’* p=0.1588), but not short-access sessions (*‘Esc4’ vs.* baseline *p=0.944, ‘Esc4’ vs.‘Esc1’* p=0.9752, *‘Esc4’ vs.‘Esc2’* p=0.9714 and ‘*Esc4’ vs.‘Esc3’* p=0.9749). Comparisons between the two groups revealed that cocaine intake was similar during baseline (Sidak, p>0.9999), while rats in long-access took more drug than animals in short-access during *‘Esc2’* (p=0.016)*, ‘Esc3’* (p<0.0001) and *‘Esc4’* (p<0.0001)

### Peer’s presence led to reduced cocaine intake, even after extended-access escalation

***In male rats maintained in short-access*** (**Fig.1D**, REML, social conditions: F(3,18)=7.311, p=0.0021, sessions: F(4,24)=0.2623, p=0.8992, interaction: F(12,68)=0.9488, p=0.5052, short-access EYFP-control group n=7) the *‘Without peer’ condition* (31.8±2.3) did not modulate cocaine intake compared to baseline (Tukey: p=0.8659), or *‘Esc4’ (p=0.97985)*). In contrast, cocaine intake was reduced in the *‘With peer’* condition (22.2±2.9 injections) compared to baseline (p=0.0153), *‘Esc4’* (p=0.008) and *‘Without peer’* (p=0.0160).

***In animals subjected to the long-access procedure*** (**Fig.1E**, REML, social condition F(3,21)=23.8, p<0.0001, session: F(4,28)=0.5061, p=0.7316, interaction: F(12,81)=0.5771, p=0.8545, long-access EYFP-control group n=8), cocaine intake ‘*Without peer*’ (53.9 ±2.7 injections) was maintained at an escalated-like level (Tukey: p=0.9998 *vs. Esc4*), rats consuming significantly more drug than baseline (p<0.0001). In contrast, drug consumption in the ‘*With peer*’ condition (34.8±3.8 injections) was decreased compared to ‘*Esc4*’ (p=0.0002), and to *‘Without peer’* (p=0.0001), reaching a level similar to baseline (p=0.6882).

Finally, the amount of cocaine taken during *‘Without peer’* and *‘With peer’* conditions was compared between male rats subjected to short-*vs.* long-access to the drug (**Fig.1F**, REML, social condition: F(3,39)=23.43, p<0.0001, access duration to the drug: F(1,13)=36.03, p<0.0001, interaction: (F(3,39)=10.53, p<0.0001, long-access EYFP-control group n=8 and short-access EYFP-control group n=7). The beneficial effect of social presence was found in both short and long-access groups, but animals maintained in short-access consumed less cocaine than rats subjected to the long-access in both ‘*Without peer’* (Sidak, p<0.0001), *‘With peer’* (p=0.0046) and *‘Esc4’* (p<0.0001) conditions.

### The STN photo-inhibition outweighed social influence and thus differently modulated the beneficial effect of the peer’s presence after limited or escalated cocaine consumption

***Before activation of the laser,*** all rats (EYFP-control (n=15) and ARCHT3.0 (n=15)) subjected to cocaine short-(n= 7 and 6 respectively for EYFP-controls and ARCHT3.0) and long-access (n=8 and 9 respectively for EYFP-control and ARCHT3.0)) took a similar and stable amount of drug during baseline (**Fig.2A**; REML, **Table 2**). Across the following short- or long-access to cocaine, REML analysis (**Table 3**) revealed significant effects of sessions (F(19,451)=8.514, p<0.0001), drug access duration (F(1,26)=61.34, p<0.0001) and their interaction (F(19, 451)=4.148, p<0.0001) on cocaine use, but no effect of the optogenetic groups (F(1, 26)=0.07892, p=0.781), while the laser was not yet activated. The amount of cocaine taken during the entire session (6h) was then averaged per blocks of 5 sessions (*‘Esc1’*, *‘Esc2’*, *‘Esc3’*, *‘Esc4’*) and compared between EYFP-control and ARCHT3.0 rats in long-access (**Fig.2B**; REML, 5-sessions block: F(3,45)=17.76, p<0.0001, optogenetic group: F(1,15)=0.03308, p=0.8581, interaction: F(3,45)=0.4097, p=0.7468, long-access EYFP-control group n=8 and long-access ARCHT3.0 group n=9). Both EYFP-control and ARCHT3.0 rats escalated their cocaine intake, reaching 139.3±5.3 injections during *‘Esc4’* (Dunnett: *‘Esc4’ vs. ‘Esc1* p<0.0001*, ‘Esc2’* p=0.0011 and*‘Esc3’* p=0.0401).

**Figure 2.**
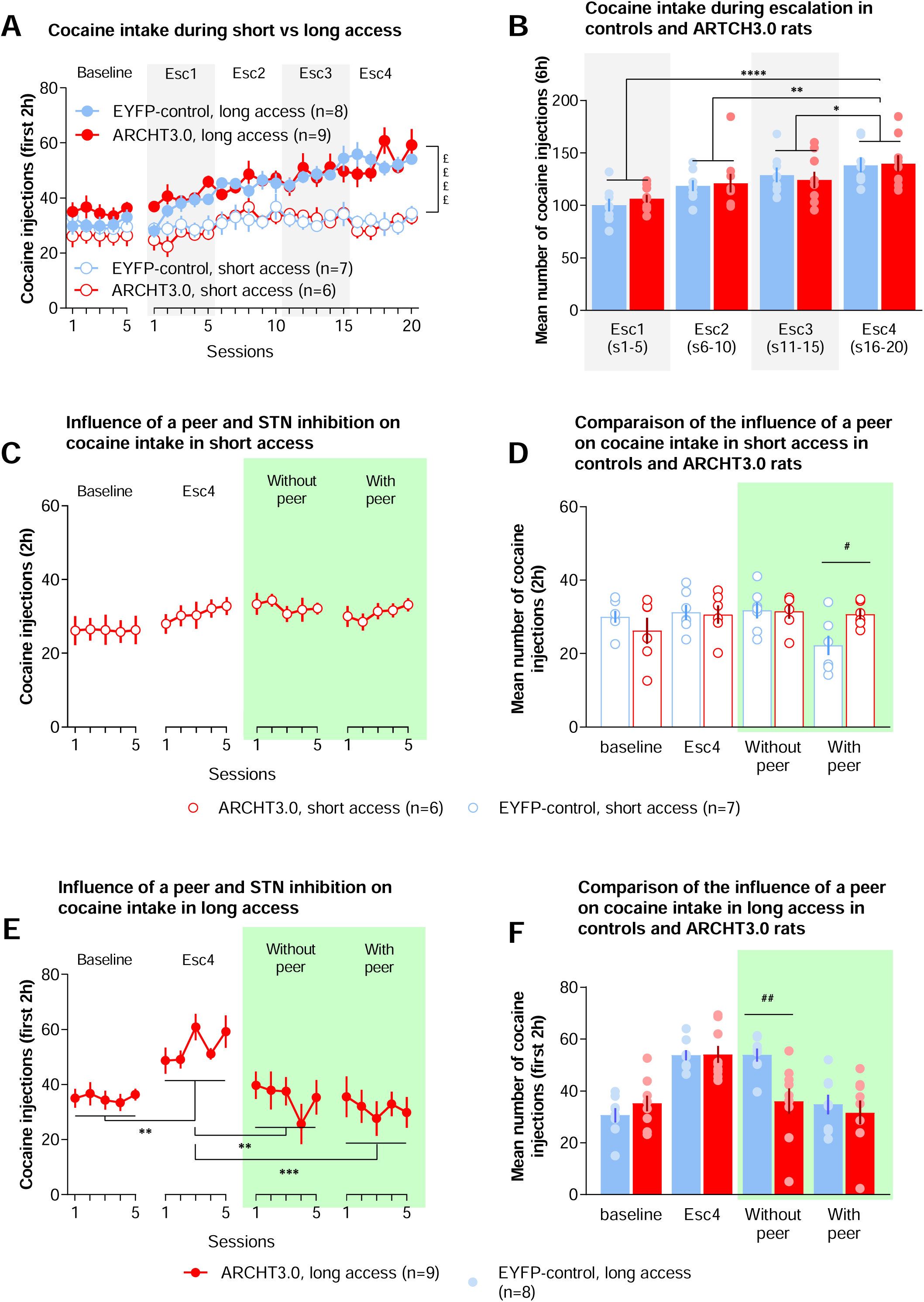
The photo-inhibition of the subthalamic nucleus mimics peer presence by reducing escalated drug consumption. **(A)** Number of cocaine injections received during the two first hours of the baseline and the twenty sessions with long (6h; n=8 EYFP-control, plain blue dots; n=9 ARCHT3.0, plain red dots) or short (2h; n=7 EYFP-control, empty blue dots; n=6 ARCHT3.0, empty red dots) access to the drug. No laser was activated during this phase. **(B)** Mean number of cocaine injections received during the six hours averaged in blocks of 5-sessions (escalation: *‘Esc1’*, *‘Esc2’*, *‘Esc3’* and *‘Esc4’*, corresponding respectively to sessions 1-5, 6-10, 11-15 and 16-20) in EYFP-control (blue bars; mean for each bar: 100.3 (±6,5), 118.7 (±5.5), 129.0 (±7.4) and 138.4 (±7.6)), and ARCHT3.0 (red bars; mean for each bar: 106.4 (±4.1), 121.2 (±9.4), 124.2 (±8.2) and 140.1 (±8.3)) in rats subjected to the long access procedure. Each dot represents an individual performance. **(C)** Cocaine injections received during the two hours of the baseline, *‘Esc4’*, *‘Without peer’* and *‘With peer’* sessions, in ARCHT3.0 rats subjected to the short access procedure. The activation of the laser is represented in green shaded area. **(D)** Mean number of cocaine injections received during the baseline, Esc4, *‘Without peer’* and *‘With peer’* conditions in EYFP-control (empty blue bars) and ARCHT3.0 (empty red bars; 26.2 (±3.9), 30.9 (±2.6), 31.9 (±1.8) and 30.8 (±1.4)) rats maintained in short access. **(E)** Cocaine injections received during the 2h-sessions of the baseline, the two first hours of *‘Esc4’ sessions*, and the 2h-sessions *‘Without peer’* and *‘With peer’* in ARCHT3.0 rats subjected to the long access procedure. **(F)** Mean number of cocaine injections received during the baseline, Esc4 (two first hours), *‘Without peer’* and *‘With peer’* conditions in EYFP-control (plain blue bars) and ARCHT3.0 (plain red bars; 35.2 (±2.9), 54.1 (±3.5), 36 (±4.7) and 31.6 (±4.4)) rats subjected to the long access procedure. *££££: p<0.0001 sessions x long vs. short access groups * p<0.05; ** p<0.01; *** p<0.001; **** p<0.0001 between conditions. #: p<0.05; ## p<0.01 between EYFP-control and ARCHT3.0 rats*.

**Table 3.**
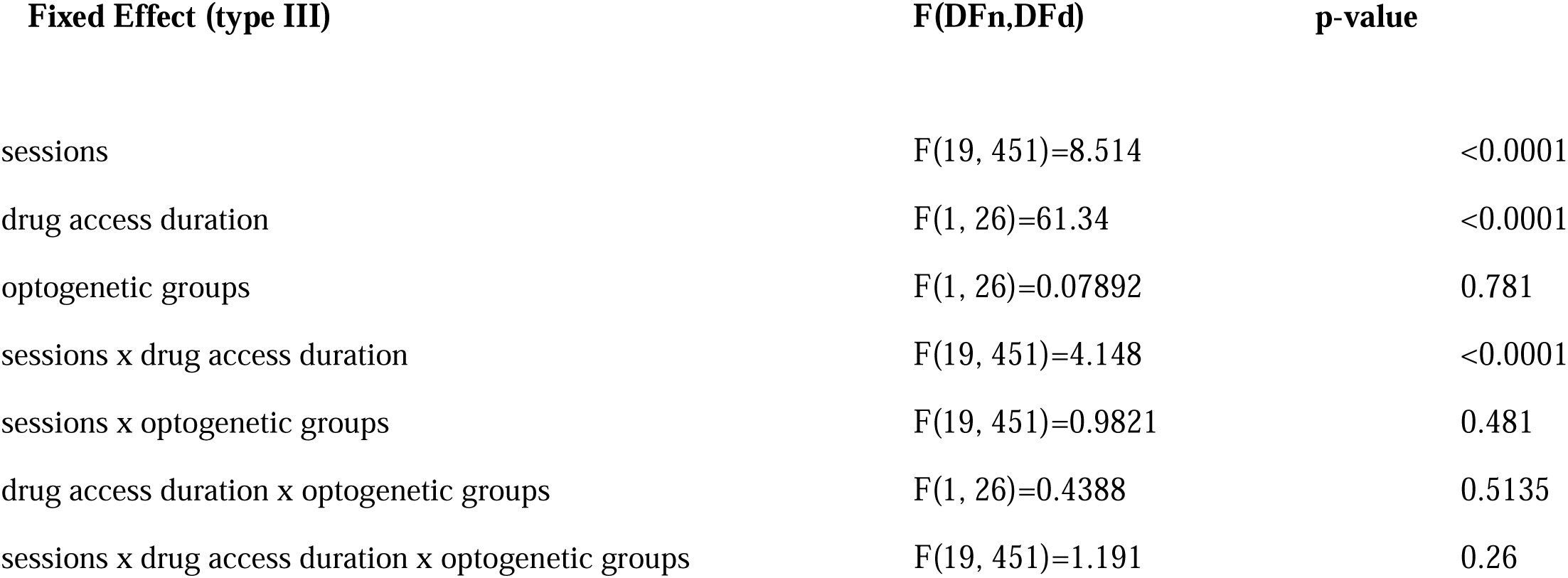
Results of the REML performed on the 2 first hours of cocaine self-administration during the twenty sessions of cocaine escalation in EYFP-control and ArchT3.0 animals subjected to short and long access to the drug.

| Fixed Effect (type III) | F(DFn,DFd) | p-value |
| --- | --- | --- |
| sessions | F(19, 451)=8.514 | <0.0001 |
| drug access duration | F(1, 26)=61.34 | <0.0001 |
| optogenetic groups | F(1, 26)=0.07892 | 0.781 |
| sessions x drug access duration | F(19, 451)=4.148 | <0.0001 |
| sessions x optogenetic groups | F(19, 451)=0.9821 | 0.481 |
| drug access duration x optogenetic groups | F(1, 26)=0.4388 | 0.5135 |
| sessions x drug access duration x optogenetic groups | F(19, 451)=1.191 | 0.26 |

***In ARCHT3.0 rats subjected to short-access*** (**Fig.2C**, REML, condition: F(3,15)=3.001, p=0.0637, session: F(4,20)=0.8369, p=0.5178, interaction: F(12,54)=0.439, p=0.94; Short access ARCHT3.0 n=6), the laser activation did not modulate cocaine intake in the *‘Without peer*’ condition, rats consuming similar amount of drug (31.9±1.8 injections) compared to baseline (26.3±3.9) and *‘Esc4’* (30.9±2.6). In the *‘With peer’* condition, cocaine consumption (30.8±1.4) was also similar to that of other conditions. This suggests that laser activation outweighed the social influence and suppressed the reduction in drug consumption induced by the peer’s presence. We then compared the cocaine intake during baseline*, ‘Esc4’*, *‘Without peer’* and *‘With peer’* between ARCHT3.0 and EYFP-control rats subjected to short-access (**Fig.2D**; REML, conditions: F(3,33)=5.131, optogenetic group: F(1,11)=0.1957, p=0.6668, p=0.0051, interaction: F(3,33)=5.656, p=0.0031; short-access EYFP-control group n=7 and short-access ARCHT3.0 group n=6). While both groups took similar amount of drug during baseline (Sidak, p=0.6725), *‘Esc4’* (p>0.9999) and *’Without peer’* (p=0.9999), ARCHT3.0 animals took more cocaine than EYFP-control ones ‘*With peer’* (p=0.0417), suggesting that the STN photo-inhibition did not affect limited cocaine intake by itself, but prevented the beneficial influence of the peer’s presence on drug consumption.

***In ARCHT3.0 rats subjected to long-access* procedure** (**Fig.2E**, REML, conditions: F(3,24)=9.912, p=0.0002, sessions: F(4,32)=1.082, p=0.382, interaction: F(12,89)=1.609, p=0.1032, long-access ARCHT3.0 group n=9), laser activation reduced cocaine intake, rats consuming less drug in the *‘Without peer’* condition than during *‘Esc4’* (two first hours; 54.1±3.5 injections, Tukey, p=0.0023), returning to baseline level (35.2±2.9 injections, p=0.9999). In the condition *‘With peer’*, ARCHT3.0 rats’ consumption (31.6±4.9) was also reduced compared to *‘Esc4’* (p=0.0003), and similar to the baseline (p=0.8604), with no additive effect compared to the *‘Without peer’* condition (p=0.8304), suggesting again that STN photoinhibition effect outweighed the influence of the social context.

The amount of cocaine taken during the baseline, *‘Esc4’* (two first hours), *‘Without peer’* and *‘With peer’* was then compared between ARCHT3.0 and EYFP-control rats subjected to the long-access procedure (**Fig.2F**; REML, optogenetic group: F(1,15)=1.07, p=0.3173, condition: F(3,45)=28.78, p<0.0001, interaction: F(3,45)=6.616, p=0.0008; long-access EYFP-control group n=8 and long-access ARCHT3.0 group n=9). Both groups took similar amount of cocaine when the laser was not activated (Sidak, baseline p=0.8491 and *‘Esc4’* p=0.9999). When the laser was activated, in *‘Without peer’* condition, ARCHT3.0 rats took less cocaine than EYFP-control animals (p=0.0035), whereas in *‘With peer’* condition, their consumption was similar (p=0.963), confirming the lack of additivity of the STN inhibition with the social influence.

Finally, the number of cocaine injections taken during baseline, *‘Esc4’* (2 first hours), *‘Without peer’* and *‘With peer’* was compared between ARCHT3.0 animals subjected to the short- and the long-access procedures (**Fig.S2**; REML, access duration: F(1,13)=4.359, p=0.0571, condition: F(3,39)=9.414, p<0.0001, interaction: F(3,39)=7.681, p=0.0004; Long access ARCHT3.0 group n=9 and Short access ARCHT3.0 group n=6). Both groups showed a similar baseline of cocaine intake (Sidak, p=0.3567). Rats subjected to the long-access procedure took more cocaine during *‘Esc4’* than animals maintained in short-access (p=0.0003). However, contrary to the EYFP-control rats, both ARCHT3.0 animals groups took similar amount of cocaine ‘*Without peer*’ (p=0.9092) and ‘*With peer*’ (p=0.9998).

### Optogenetic STN HF-stimulation produced similar results as optogenetic inhibition

hChR2 rats (n=9, **Fig.3A**) took 30±4.7 cocaine injections during baseline, a level similar to that of the EYFP-control later subjected to the long-access (**Fig.3B**, detailed results in supplemental). They also escalated their cocaine consumption during the two first hours of the 20 long-access sessions, reaching 63.5±7.4 injections at the last session. The total number of cocaine injections taken during the 6h-sessions across *‘Esc1’*, *‘Esc2’*, *‘Esc3’*, *‘Esc4’*) by hChR2 rats was similar to EYFP-control rats (**Fig.3C**). hChR2 rats reached a consumption of 135.9±9.1 injections during *‘Esc4’ (vs. ‘Esc1’*, Dunnett, p<0.0001). During the two first hours of the sessions, with laser activated, hChR2 rats consumed less cocaine in the *‘Without peer’* condition (35.4±2.9 injections) than during *‘Esc4’* (55.2±4.7, Tukey, p=0.0066), returning to baseline level of drug use (30.0±4.7 injections, p=0.7227) (**Fig.3D**) as it was for the ARCHT3.0. Their cocaine consumption was not further reduced in the *‘With peer’* condition (25.1±4.8 injections, Tukey, p=0.2222),that was equivalent to the baseline (p=0.786) and significantly lower than *‘Esc4’* (p<0.0001), suggesting that STN photo HF-stimulation also outweighed the influence of the social context.

**Figure 3.**
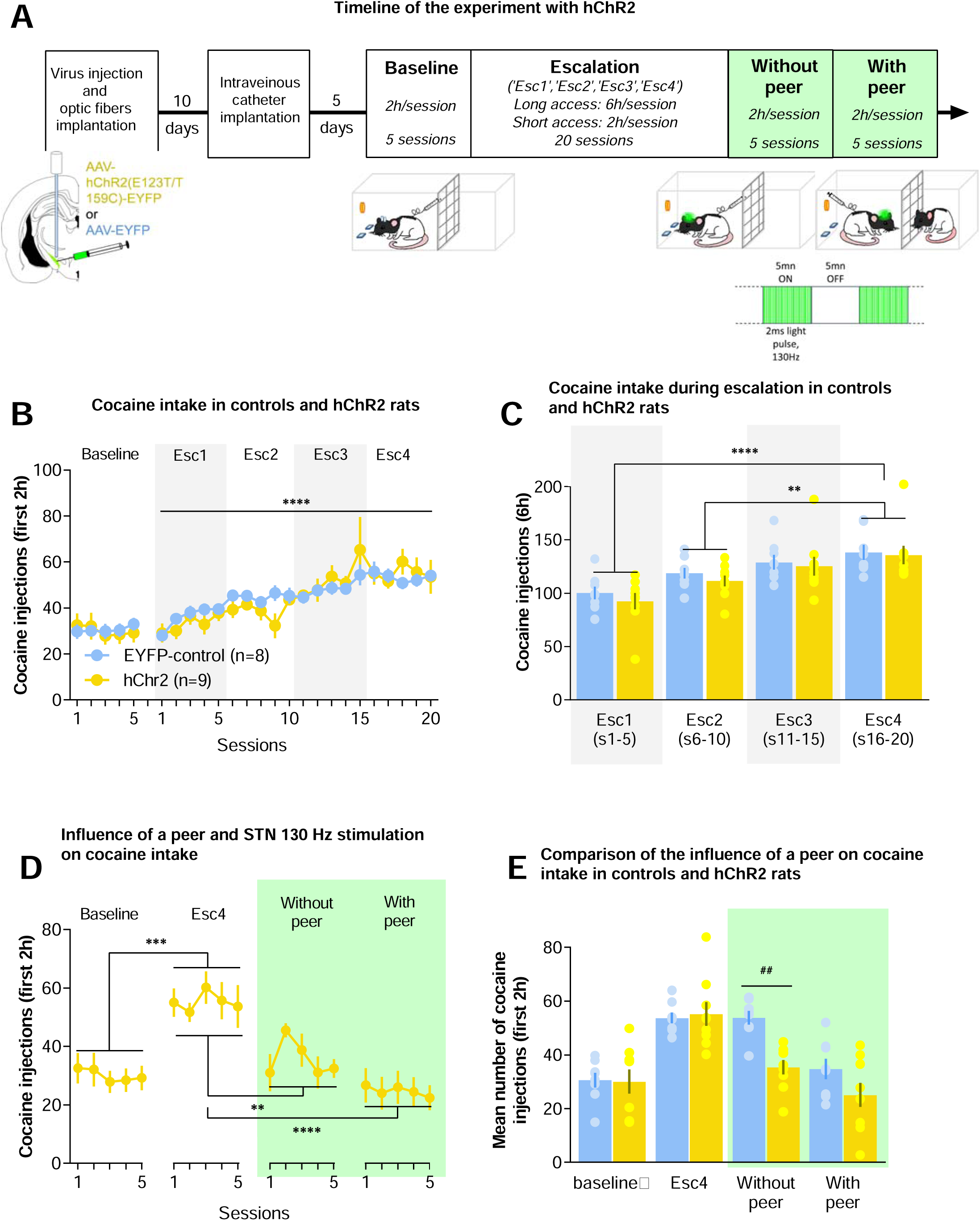
The high frequency photo-stimulation of the subthalamic nucleus mimics peer presence at reducing escalated drug consumption. **A**) Timeline of the experiment with hChr2 **(B)** Number of cocaine injections received during the two first hours of the baseline and the twenty sessions with long access to the drug (n=8 EYFP-control, plain blue dots; n=9 hChR2, plain yellow dots). (**C**) Mean number of cocaine injections received during the six hours averaged in blocks of 5-sessions (escalation: ‘Esc1’, ‘Esc2’, ‘Esc3’ and ‘Esc4’, corresponding respectively to sessions 1-5, 6-10, 11-15 and 16-20) in EYFP-control (blue bars; mean for each bar: 100.3 ±6.5, 118.7 ±5.5, 129.0 ±7.4 and 138.4 ±7.6), and hChR2 rats (yellow bars; mean for each bar: 92.6 ±8.1, 111.6 ±5.5, 125.4 ±9.3 and 135.9 ±9.1). Each dot represents an individual performance. No laser was activated during this phase. (**D**) Cocaine injections received during the two hours of the baseline, the two first hours of ‘Esc4’, and the 2h-sessions ‘*Without peer*’ and ‘*With* peer’ in hChR2 rats. (**E**) Mean number of cocaine injections received during the baseline, Esc4 (two first hours), *‘Without peer’* and ‘*With peer*’ conditions in EYFPcontrol (plain blue bars) and hChR2 (plain yellow bars; 30.0 ±4.7, 55.2 ±4.7, 35.4 ±2.9 and 25.1 ±4.8) rats. *\*\* p<0.01; *** p<0.001; **** p<0.0001 between conditions. ## p<0.01 between EYFP-control and hChR2 rats.*

The amount of cocaine taken during baseline, ‘*Esc4*’ (the two first hours), *‘Without peer’* and *‘With peer’* was then compared between hChR2 and EYFP-control rats in long-access procedure (**Fig.3E**). Both groups showed similar cocaine intake when the laser was not activated (Tukey, baseline p>0.9999 and *‘Esc4’ p=0.9972)*. With laser activation aiming at STN HF-stimulation, while *‘Without peer’* hChR2 animals took less drug than EYFP-control (p=0.0024), in the *‘With peer’* condition, cocaine consumption was similar between groups (p=0.2229).

### Electrophysiological assessment of STN photo-inhibition

STN neurons electrophysiological properties, including resting membrane potential (Kruskall-Wallis: p=0.8214), cell capacitance (p=0.0644), membrane resistance (p=0.7397), access resistance (p=0.3158) and firing rate (mixed 2way ANOVA, Frequency x Group interaction: F(30,390)=0.2480, p>0.9999) were equivalent between EYFP-control (n=3 animals; n=8 cells) and ARCHT3.0 (n=7 animals; n=11 cells) (**Fig.S3**). Efficacy of the optogenetic modulation was evaluated by applying light pulses (**Fig.S4**). Longer pulse-width and higher light intensity elicited higher inhibitory currents in ARCHT3.0 STN neurons (at 5mW, pulse duration; χ²(5)=51.88, p<0.0001, light intensity: χ²(3)=22.00, p<0.0001). Consistent with previous reports (41), higher laser activation (*i.e.,* increased pulse duration and brighter illumination) leads to higher activation of opsins and increases the transmission through a larger volume of tissue, thereby recruiting a larger pool of neurons. To further assess the effect of optogenetic inhibition, electrical stimuli (10Hz, injected current: rheobase + 20-130pA) were delivered to STN neurons, to evoke action potentials before applying photo-inhibition (**Fig.S5A**). A 15s light-pulse delivered to ARCHT3.0-expressing neurons successfully inhibited neuronal firing (**Fig.S5B**, Dunn, p=0.0006). We then applied the same light pattern than during the behavioral experiments (*i.e.,* alternation of a 15s period ON and 5s period OFF, for 5min) on other ARCHT3.0 transfected cells (n=2 rats, n=7 cells) (**Fig.S5C**). Photo-inhibition of STN neurons leads to reduced firing rate (**Fig.S5D**; Dunn, p=0.0323), which resumed when the laser was stopped (p=0.9999). Thus, our photo-inhibition protocols efficiently inhibited STN neurons *ex vivo*.

STN photo-inhibition was then tested *in vivo* by performing STN extracellular recordings in ARCHT3.0 anesthetized animals (n=4). A 15s light-pulse (5mW) successfully inhibited STN firing (**Fig.S5E,F**, Dunn, p<0.0001, n=16 cells) but did not impact neuronal activity during the intermittent 5s OFF periods (p=0.959), as in our *ex vivo* preparation.

Accordingly, the normalized firing frequency of ARCHT3.0-transfected neurons significantly diminished during the 180s laser bins (alternating 15s ON/5s OFF periods) compared to baseline (**Fig.S5G**; Dunn, p<0.0001). STN firing during the 20s post-laser period was similar to the baseline activity (p>0.9999).

### Electrophysiological assessment of STN HF photo-stimulation

Our behavioral results showed that STN HF photo-stimulation mimics the effects of the STN photoinhibition. To check whether HF photo-stimulation of the STN has a different effect than its photoinhibition on the neuronal activity, we performed both *ex vivo* and *in vivo* recordings. Naïve adult animals were injected with AAV5-CaMKII-hChR2 (E123T/T159C)-p2A-EYFP-WPRE (n=14) and recorded 3-4 weeks post-injection.

*Ex vivo* electrophysiological properties of hChR2-transfected STN neurons (n=4 animals, n=10 cells) were similar to those of EYFP-control and ARCHT3.0 transfected cells (**Fig.S3**). Discrete light pulses were applied in hChR2expressing STN neurons (n=10) (**Fig.S5D-G**). Longer pulse-width and higher light intensity elicited higher depolarizing currents (intensity 10mW, pulse duration: χ²(5)=46.00, p<0.0001, light intensity: χ²(3)=28.92, p<0.0001). Optogenetic stimulation effects were assessed with 10s laser illumination at various frequencies (**Fig.S6A**). Higher frequencies of stimulation did not change the number of action potentials compared to lower frequencies (**Fig.S6B**). However, higher frequencies significantly increased membrane potential (**Fig.S6C**), suggesting an increase of the STN neurons excitability. To test this hypothesis, a 10Hz electrical stimulation was applied in other hChR2-transfected STN neurons (n=2 rats, n=8 cells) (injected current: Rheobase +0-50pA) to evoke few action potentials. We then applied 130Hz laser stimulation for 5min (**Fig.S6D**), which significantly increased the firing rate of STN neurons (Dunn, p<0.0351, **Fig.S6E**). Following cessation of the photo-stimulation, the firing rate was no longer different from the baseline (Dunn, p>0.05). This suggests that 130Hz optogenetic stimulation is unlikely to induce action potentials on its own but increases STN neurons excitability and does not result in a neuronal inhibition.

The impact of STN HF-stimulation (130 Hz during 3 min) was also assessed *in vivo* in hChR2-transfected rats (n=7 animals; n=16 cells). The population response was estimated by normalizing the STN firing using 1-min baseline prior the photo-modulation. Although the overall population increased its firing rate during light stimulation (**Fig.S6F, G**), about half of the cells (n=7) did not modify their firing rate (non-responding cells; Dunn, p=0.3629). In contrast, the impact of STN HF photo-stimulation was strongly marked in 9 neurons (responding cells; Dunn, p=0.0019). These cells returned to their basal activity when the photo-stimulation was turned OFF (p>0.9999). Thus, STN HF photo-stimulation does not inhibit neuronal activity, but rather increases it.

## Discussion

Our study reveals that the presence of a non-familiar, cocaine-naïve, peer reduced both limited and escalated cocaine intake in male rats, while having no effect on limited intake in females. STN photo-inhibition reduced escalated, but not limited, cocaine use. When combined with peer presence, STN optogenetic inhibition outweighed the social influence, preventing the beneficial effect under short-access conditions and blocking a potential additive effect under long-access conditions. STN HF photo-stimulation mirrored the effects of photo-inhibition on escalated cocaine consumption.

### STN Photo-Modulations Reverse Cocaine Escalation Without Altering Limited Intake

Extended drug access led to cocaine intake escalation, reflecting the compulsive-like loss of control seen in humans (39) and inducing compulsive-like drug-seeking behavior in rats, unlike short access (34). Compared to short-access animals, long-access rats maintained higher cocaine intake even when session durations were reduced from 6 to 2 hours, consistent with previous findings (42)

Both STN photo-inhibition and HF-stimulation nearly halved cocaine consumption post-escalation. Laser activation had no effect in EYFP-control animals, revealing no side effects such as potential tissue heating due to prolonged laser exposure (43).Consistent with previous reports using other inhibition methods (29–31), STN photo-inhibition, validated by electrophysiological experiments, decreased escalated but not limited cocaine intake. This differential effect may be linked to earlier findings that STN inhibition does not affect cocaine intake under an FR1 reinforcement schedule (considered a “no effort” task) but reduces motivation in a progressive ratio schedule requiring increased effort to obtain the drug (29). After extended drug access, animals typically show heightened motivation for the drug (44), accepting effort levels previously deemed excessive, likely reflecting a shift in the subjective valuation of reward versus effort. The STN has been implicated in encoding both reward value and effort (45–48). Extended drug access may alter this encoding in the STN, explaining why STN photo-inhibition is effective only after a loss of control over cocaine use in the FR1 paradigm.

One limitation of this study is that only a single dose of cocaine was tested, potentially masking sensitivity shifts induced by STN manipulation. However, previous findings demonstrated that STN inhibition induces a downward vertical shift, not a lateral one, during dose-response testing (29).

Past studies have shown that STN HF-DBS reduces cocaine intake post-escalation when applied after a period of abstinence (31). In contrast, STN photo-modulations in this study immediately reversed cocaine escalation. Optogenetic manipulations may outperform electric HF-DBS (49), potentially due to their greater specificity compared to electric stimulation or chemical lesions, which can affect neighboring structures and passing fibers. This enhanced specificity may better suppress STN low-frequency oscillatory activity associated with cocaine escalation (31,34).

Although it may seem counterintuitive that STN photo-inhibition and HF photo-stimulation yielded similar behavioral effects, this finding aligns with previous results showing that chemically-induced inhibition of the STN and its HF electric stimulation both reduce cocaine-related motivation (29,30). This was attributed to HF-DBS simultaneously inactivating cell bodies while stimulating passing fibers. Our STN electrophysiological recordings confirmed proper inhibition of STN neuron activity in ARCHT3.0 animals. However, STN optogenetic HF-stimulation resulted in either no effect or neuronal activation, contrasting with the transient inhibition induced by electric STN HF-DBS (50). This discrepancy does not fully explain the similar behavioral outcomes observed. One hypothesis is that modulating STN activity, regardless of direction, may suppress abnormal synchronized oscillatory activity induced by extended drug access. Both optogenetic HF-stimulation and inhibition have been shown to suppress pathological beta oscillations in computational models of parkinsonian neural activity (49). In another study on 6-OHDA parkinsonian rats, 130 Hz STN photo-stimulation increased the firing rate in 50% of neurons but decreased it in 32%, potentially due to recruitment of inhibitory neurons (51). This inhibitory effect may contribute to the similar behavioral outcomes observed with photo-inhibition and HF-stimulation in this study. Alternatively, repetitive high frequency optogenetic stimulation of ChETA-transfected amygdala inputs to the prefrontal cortex was shown to induce a long-lasting synaptic depression (52), suggesting that high frequency photo-activation could indeed evoke inhibitory responses. A detailed analysis of the behavior during the laser bins (*i.e.,* 5min ON and OFF periods) was not possible in our operant chambers. Such analysis could have allowed us to better understand why similar behavioral effect could be observed with the 2 opsins despite their opposite effects on STN neuronal activity.

Further electrophysiological recordings of STN efferent structures are necessary to clarify how STN HF-stimulation could evoke inhibitory responses. Nonetheless, from a translational perspective, our findings confirm STN neuromodulation as a promising treatment for substance use disorder (SUD), as previously suggested (29–35,53).

### Beneficial Effect of Peer Presence

In females, the presence of a cocaine-naïve unfamiliar congener, changed daily, did not influence cocaine intake under short-access conditions. Female rats are generally less sensitive to social interactions than males (54) and exhibit weaker social preference over cocaine (17). This aligns with findings showing that females, unlike males, do not reduce cocaine intake when housed with a cocaine-naïve peer separated by a grid and absent during drug consumption (55). Interestingly, both sexes consume more cocaine when housed with a cocaine-using peer (22,55). The absence of a protective social effect in females suggests greater sensitivity to detrimental rather than protective social influences. Additionally, female rats acquire (56) and escalate (57) drug self-administration more rapidly than males. This points to sex-dependent mechanisms underlying drug consumption, potentially contributing to differences in social context sensitivity during drug use. Future studies should examine further the influence of the presence of a female peer also on drug intake and the role of STN.

In males, our findings expand upon previous results showing that a drug-naïve observer reduces cocaine intake during short access (25) and escalation procedures under different conditions (27). We demonstrated a similar beneficial effect following cocaine escalation. Importantly, the stimulus peer was a daily-changed unfamiliar to prevent habituation, as familiarity diminishes social influence on drug intake (25,26). The use of unfamiliar peers maintained highly rewarding social novelty (25,54).

Notably, the beneficial influence of peer presence on cocaine intake in males appears independent of cocaine dose, as similar effects were observed in this study (250 µg cocaine/injection) and previous work (80 µg cocaine/injection) (25).

Our findings suggest that the presence of a drug-naive peer can protect against SUD, supporting the wider implementation of consumption rooms, where drug users can consume substances in a medically supervised, socially supportive environment. A recent report highlighted reduced opioid overdose deaths due to such facilities (58).

However, peer presence does not always protect against drug use. When rats learn drug self-administration together, peer presence facilitates acquisition (22,24,59). Similarly, when cocaine availability is paired with a peer’s presence, this association can later trigger drug craving and relapse (60). Altogether, these findings highlight that peer influence on drug intake depends on factors such as social contact contingencies, sex (22,55), peer behavior (abstinent or drug user) (22,24,25,27,61), relationship with the focal rat (e.g., familiarity, dominance) (25,26), and drug type (62). As others have noted (5,63), incorporating social context into SUD animal models is crucial for addressing the translational crisis in addiction research and developing effective treatments for SUD.

### Social Influence Relies on the STN Network

Peer presence significantly reduced cocaine intake under both short-(below baseline) and long-access (to baseline) conditions in control males. In short-access conditions, STN photo-inhibition abolished the peer’s beneficial effect, indicating a direct causal role for the STN in social modulation. In long-access rats, STN photo-inhibition or HF-stimulation alone reduced drug consumption. However, while peer presence reduced cocaine intake in the EYFP-control group, no additive effect was observed with concurrent STN photo-modulation. In both conditions, STN modulation overrode social influence, resulting in equivalent cocaine intake regardless of peer presence. This finding is consistent with prior studies showing that STN lesions prevented the protective effect of positive ultrasonic vocalizations on limited cocaine intake, despite having no effect independently (26,28). These results indicate that an intact STN is necessary for social modulation of drug use. Moreover, an fMRI study demonstrated STN involvement in modulating social influence on stop-signal reaction times in human cocaine users (64).

We recently showed that STN HF-DBS suppressed peer-induced increases in alcohol consumption (62), further confirming that the STN modulates peer influence on drug intake, regardless of the drug type or social effect (increase or decrease in drug use).

## Conclusion

While the presence of a drug-naïve unfamiliar congener had no effect on limited intake in females, it decreased both limited and addiction-like cocaine consumption in males. Optogenetic inhibition of the STN, as well as its HF photo-stimulation, reduced addiction-like but not limited drug use, confirming the potential of STN HF-DBS in treating cocaine addiction. When combined with social presence, STN manipulation exerted an overriding influence. These findings highlight the crucial role of the STN in mediating the beneficial effects of peer presence on drug consumption. Further research is needed to elucidate the involvement of the STN in social modulation of drug use and the underlying cellular mechanisms.

## Declaration of generative AI and AI-assisted technologies in the writing process

During the preparation of this work the authors used ChatGPT in order to correct the English in abstract, introduction and discussion. After using this tool/service, the author(s) reviewed and edited the content as needed and take full responsibility for the content of the publication.

## Data availability

All data reported in this paper will be made available by the lead contact upon reasonable request.

## Funding sources

Centre National de la Recherche Scientifique (CBa, NM, MD, CBr)

Aix-Marseille Université (ATC, CBa, CV, LV, FP, MN, MD, YP)

French Ministry of Higher Education, Research and Innovation (ATC, CV)

Mission Interministérielle pour la Lutte contre les drogues et les conduites addictives, Programme Apprentis Chercheurs MAAD (CBa)

Fondation pour la Recherche Médicale (FRM DPA20140629789) (CBa) Institut de Recherche en Santé Public (IRESP-19-ADDICTIONS-02) (CBa)

Institut National Du Cancer (French National Cancer Institute) (INCA-16032) (LV) Agence Nationale de la Recherche (ANR-21-CE16-0002-01) (MD)

None of these funding sources had no further role in study design; in the collection, analysis and interpretation of data; in the writing of the report; and in the decision to submit the paper for publication.

## Author contributions

CV, ATC, YP, and CBa conceptualized and designed the study. CV, LV and JM, performed the behavioral experiments, and CV and LV analyzed them. CV, LV, FP and CBr performed immunohistochemistry and histology. ATC performed and analyzed the in-vitro electrophysiological recordings. NM performed the in-vivo electrophysiological recordings and their analyses. CV, ATC, and CBa wrote the original draft. CV, LV, NM, JM, MD, YP, and CBa reviewed and edited the original draft.

## Acknowledgments

The authors thank Dr G F Koob for critical reading of the manuscript, J Baurberg for technical assistance with all electronic devices, Drs. F Brocard and R Bos for allowing use of their patch clamp equipment, and the technical staff of the animal facility for ensuring the well-being of our animals.

