## Supplementary data for "Presence of an unfamiliar peer reverses escalated cocaine intake in male rats: involvement of the subthalamic nucleus"

**Supplementary Materials and Methods:**

### Animals

Lister Hooded rats (~320 g upon arrival) (Charles River Laboratories, Saint-Germain-surl’Arbresle, France) were subjected to behavioral experiments (n=50 males and n=8 females), or used as observer peers or for electrophysiological recordings (n=30 males and n=5 females). Animals were pair-housed and provided *ad libitum* access to water and food (Scientific Animal Food and Engineering, Augy, France), maintained in a temperature-controlled room, under a 12hour inverted light/dark cycle, with experiments conducted during the dark cycle (7am-7pm), and daily handled. All animal care and use adhered to French regulation (Decree 2010-118) and was approved by the local ethics committee and the French Ministry of Agriculture under license #03129.01 and followed the 3R European rules. Group sizes were determined considering the 3Rs principles to minimize animal usage and the targeted success rate for STN localization in our lab.

### Viruses

To transfect STN neurons, we used an AAV5 with the recombinant protein expression under CamKIIapromoter control (UNC Vector Core, Chapel Hill, USA). The EYFP-control, ARCHT3.0 and hChR2 groups received respectively the following AAV5 constructs: AAV5-CaMKII-EYFP, AAV5-CaMKII-

ArchT3.0-p2A-EYFP-WPRE and AAV5-CaMKII-hChR2(E123T/T159C)-p2A-EYFP-WPRE.

### Surgeries

Males Rats weighing ~400g were anesthetized with ketamine (Imalgene, Merial, 100 mg/kg, *i.p*.) and medetomidine (Domitor, Janssen 30 mg/kg, *i.p.*), reversed with atipamezole (Antisedan, Janssen, 0.15 mg/kg, *i.m*.) at the end of the surgery. They received amoxicillin (Duphamox, LA, Pfizer, 100 mg/kg, *s.c*.) and meloxicam (Metacam, Boehringer Ingelheim, 1 mg/kg, *s.c.*) for antibiotic treatment and analgesia, respectively. Bilateral injections of 0.45µL AAV5 virus into the STN were performed using a stereotaxic frame (David Kopf apparatus) (with tooth bar set at -3.3 mm), at the following coordinates : anterior/posterior = -3.7 mm ; lateral = ±2.4 mm from bregma ; dorsoventral = -8.35 mm from skull; from Paxinos & Watson, (40)), averaging bregma and interaural coordinates. Optic fibers for behavioral tests were implanted 0.4 mm above each injection site within a dental cement head-cap, with screws anchored on the skull.

After a week, rats (n=50 males and n=8 females) underwent intra-jugular catheter implantation for drug self-administration. Using standard surgical procedures (29), and the same anesthetic procedure than above, silicon catheters were inserted into the right jugular vein and exited dorsally between the scapulae. Rats were then given a 10-day recovery period.

Catheters were flushed daily with a sterile saline solution containing heparin (Heparin Sodium, Sanofi, 3g/L) and gentamicine (Pangram, Virbac, 4g/l) throughout the experiment for patency maintenance and infection risk reduction. Catheter patency was also confirmed using propofol (Propovet, Abbott, 10 mg/ml) at the end of the experiment or after each session if doubts arose.

### Behavioral experiment

#### 1 - Apparatus

Homemade self-administration chambers (60x30x35cm) with two compartments separated by a metallic grid were utilized (25). The wall of one compartment per cage was equipped with two levers and a central light, which control was managed through a custom-built interface and software (developed by Y. Pelloux).

#### 2 - Experimental procedure of cocaine self-administration

The different groups of rats run the protocol in parallel (i.e. at the same time and under the same conditions). Rats’ catheters were connected to cocaine syringes on pumps (Razel Scientific Instruments, St-Albans, VT, USA), via infusion lines and liquid swivels. Under an FR1 reinforcement schedule, each press on the ‘active lever’ (counterbalanced between rats) triggered an intravenous cocaine infusion (250μg per 90μl infusion in 5s) and activated the cue-light during delivery, followed by a 20-s time-out period, during which any further active-lever press was recorded as perseveration but had no consequence. Pressing on the inactive lever was recorded but had no consequence. Only one dose of cocaine was used in these experiments, as it is classically used and has previously been used in our former work testing the influence of social context or STN manipulation (26,31,34).

Rats initially underwent 2-h daily sessions of cocaine self-administration until consumption stabilized (<25% variability in injections for 5 consecutive days) - i.e., baseline consumption. Then, females self-administered cocaine in a presence of a naïve to cocaine stranger female for 5 sessions, the abstinent congener being changed every day to prevent habituation and progressive familiarity. Males underwent either 6-h (long access groups) or 2-h (short access groups) daily sessions for 20 days. From day 10, rats' optic fibers were connected to the optic coupler during the first 2 hours of sessions, with the laser OFF, for habituation to the cable. Finally, rats underwent 10 additional 2-h sessions with laser stimulation (ON). This comprised two blocks of five consecutive days, with rats self-administering cocaine alone or with a male observer peer in the adjacent cage compartment. As for the female group, to account for potential social recognition memory effects, a different observer peer was used daily. The order of 'Without peer' vs 'With peer' conditions and the presentation of observers was counterbalanced between rats.

#### 3 - Optogenetic manipulation during behavioral testing

Homemade optic fibers (230µm, Thorlabs) were connected to a 200mW 532nm DPSS laser through an optic coupler (FCMM625-50A, Thorlabs). Light pulses, controlled by a signal generator (DS8000 Digital Stimulator, World Precision Instruments) followed specific parameters: 2ms light pulse at 130Hz pulse train for STN HF-stimulation and 15s light pulse at 0.2Hz pulse train for STN inhibition. During experiments, light pulses were discontinuous, alternating 5-min intervals of light ON with 5-min intervals of light OFF. Before experiments, light power at fiber tip was set using a power meter (PM20A, Thorlabs) to 10mW for STN HF-stimulation and 5mW for inhibition. Control rats were randomly assigned to one light stimulation condition throughout the experiment. The effects of these light delivery parameters on the STN neurons firing pattern were tested in vitro and in anesthetized animals. The choice of 130 Hz for the stimulation group aimed at mimicking the electric deep brain stimulations targeting the STN applied at 130 Hz and inducing behavioural effects that are often similar to STN lesions (30, 35). The electric stimulation applied at 130 Hz has been shown to result in STN neuronal inhibition, while stimulating passing fibers.

### Intracellular *ex vivo* recordings of STN neurons

Male rats were anesthetized with ketamine/medetomidine and perfused intra-cardiacally with ice-cold Nmethyl D-glucamine (NMDG)-based solution containing (in mM): 92 NMDG, 2.5 KCl, 1.25 NaH2PO4, 30 NaHCO3, 20 HEPES, 25 glucose, 2 thiourea, 5 Na-ascorbate, 3 Napyruvate, 0.5 CaCl2 and 10 MgSO4. The brain was then removed from the skull and glued to the stage of a vibratome (Leica, VT1000S) where it remained submerged in ice-cold oxygenated NMDG-based solution. Coronal slices (200 µm) containing the STN were collected and transferred to recover at 33°C for 10 min in a solution containing 92 mM NaCl instead of NMDG plus 1 mM MgCl and 2 mM CaCl2. Slices were then stored in normal artificial cerebrospinal fluid (ACSF) containing (in mM): 125 NaCl, 2.5 KCl, 1.25 NaHPO4, 1 MgCl, 2.4 CaCl, 26 NaHCO3 and 11 glucose. For the recordings, slices were transferred one at a time to a submersion-type chamber and perfused continuously with warm ACSF (31°C) at a rate of 3 ml/min. All solutions were continuously equilibrated with 95% O2 / 5% CO2. Neurons were visualized on an upright microscope (Olympus, BX51WI) equipped with DIC optic. Patch-clamp electrodes (4-6 MΩ) were prepared from filamented borosilicate glass capillaries (PG150T-7.5, Harvard Apparatus) using a micropipette puller (PC-10, Narishige) and were filled with an intracellular solution containing (in mM): 140 potassium gluconate, 5 NaCl, 2 MgCl2, 10 HEPES, 0.5 EGTA, 2 ATP, 0.4 GTP, pH adjusted to 7.35 and osmolarity adjusted to 270-280 mOsm/L. Series resistance was monitored during experiment and cells showing more than 20% of input resistance variation were excluded from the analysis. Recordings were made with pClamp 10.3 software using a MultiClamp 700B amplifier set with 4 kHz low-pass Bessel filter and 10 kHz digitization (Molecular Devices, Sunnydale, CA). The laser was directed at the STN using a 200μm optic fiber submerged into the bath. Laser intensities were adjusted to match those used in behavioral experiments. Cells expressing inhibitory opsins ARCHT3.0 were driven at 10Hz, with injected current intensity corresponding to rheobase + 20-130pA in order to induce stable and robust firing pattern. Cells expressing excitatory opsins hChR2 were driven at 10Hz, intensity set at +0-50pA to elicit a few action potentials in a stable pattern. Stability of the excitation was assessed trough a 2min baseline recording, then we applied the laser patterns used for behavioral experiments for 5min and the cells were monitored for another 5min with the laser OFF. For experiments testing the influence of light intensity and pulse width, cells were held at -60mV in current clamp configuration and discrete light pulses were applied. When testing the influence of light intensity, pulse duration was set to 15s and 1s for inhibition and stimulation groups, respectively. Testing the influence of the pulse duration was performed at constant light intensity: 10mW (Stimulation) and 5mW (Inhibition).

### Extracellular *in vivo* recordings of STN neuronal activity

STN optogenetic modulation was evaluated in 11 rats (n = 4, ARCHT3.0 group and n = 7, hChR2) by *in vivo* electrophysiological recordings. Ketamine/xylazine (100/10 mg/kg i.p.) anesthetized rats were mounted in a stereotaxic head frame (Horsley-Clarke; Unimécanique, Epinay-sur-Seine, France). Body temperature was maintained at 36.5°C with a homeothermic blanket controlled by a rectal probe (Harvard Apparatus, Holliston, MA). Single-unit activity in the STN was recorded extracellularly using glass micropipettes (25-35 MΩ) filled with 0.5 M sodium chloride solution. Action potentials were recorded using an Axoclamp-2B amplifier (Molecular Devices, San Jose, CA), amplified, filtered with an AC/DC amplifier (DAM 50; World Precision Instruments) and sampled at 10 kHz rate on a computer connected to a CED 1401 interface and off-line analyzed using Spike2 (Cambridge Electronic Design, Cambridge, UK). An optical fiber (230 μm-Thorlabs), inserted at a 15° angle, was lowered just above the STN and connected to a 200mW 532nm DPSS laser. The entry point coordinates were AP: -3.7 mm, ML: +1.0 mm. The fiber tip was at a depth of 7.3 mm from the cortical surface.

The STN neuronal activity was analyzed 20s before, 180s during and 20s after photo-modulation. Parameters of light stimulation matched those in behavioral experiments: ARCHT3.0-expressing rats received 15s ON/5s OFF stimulation (9 consecutive bins for 180s) at 5 mW (±10%) power and hChR2expressing rats received 130 Hz, 2 ms width stimulation at 10mW (±10%) power.

### Histology

At the end of experiments, rats were deeply anesthetized with ketamine (200 mg/kg) and medetomidine (60 mg/kg) through their catheter (behavioral testing) or intraperitoneally (in vivo electrophysiological recordings) and then perfused intra-cardiacally with 4% paraformaldehyde in PBS. Brains were extracted, cryo-protected in sucrose (30%), frozen in liquid isopentane, and sectioned in coronal slices (40µm) with a cryostat.

In hChR2 rats, native fluorescence, while present, required immuno-staining to delineate viral expression boundaries. After PBS rehydration (3 x 5 min), brain sections underwent a 90 min permeation step (PBS, 1% bovine serum albumin (BSA) 2% normal goat serum (NGS), 0.4% TritonX-100), PBS washes (3 x 5 min) and incubation with primary antibody (mouse anti-GFP, A11120, Life technologies;

1:200, in PBS 1% BSA, 2% NGS, 0.2% TritonX-100) at 4°C overnight. Sections were then washed with PBS (3 x 5 min), followed by 2h incubation at room temperature with secondary antibody (Goat antimouse Alexa 488, A11011, Life technologies, 1:400 in PBS 1% BSA, 2% NGS) and finally washed with PBS (3 x 5 min).

All brain sections were mounted on glass slides with homemade mounting medium and examined by an experimenter blinded to experimental conditions using an epifluorescence microscope (Zeiss, Imager.z2). Animals with improper fluorescence localization or optic fiber misplacement were excluded from the results (n=10 in total). Representative correct optic fibers’ placement and fluorescence expression are illustrated in **Fig. 1A**.

### Statistical analysis

All variables are expressed as mean ± SEM, statistical tests were two-tailed, with a significance threshold set at α=0.05. Statistical tests and graphs were performed using GraphPad Prism 8.0 or R (package lme4).

#### 1 - Behavioral analyses

A Grubbs test initially identified an outlier animal based on mean baseline cocaine intake in all rats (n=40 post-histological assessment). This EYFP-control rat in long access procedure took 84 (±6.3147) cocaine injections during its baseline (Grubbs test results: Z=4.152, p<0.05) and was thus excluded from statistical analysis and results.

The number of cocaine injections was analyzed by fitting linear mixed models, to account for missing data (due to catheter disconnections) at random. Were considered as fixed factors: optogenetic groups (EYFP-control, ARCHT3.0, or hChR2), access duration to cocaine (short or long access) as between factors, and sessions or conditions (baseline, escalation: ‘Esc1’, ‘Esc2’, ‘Esc3’, ‘Esc4’, ‘With peer’ and in ‘Without peer’) as within factors, and the subjects as random factor. This mixed model relies on a compound symmetry covariance matrix and is fit using Restricted Maximum Likelihood (REML). REML were followed by appropriate post hoc analyses. Effect sizes are reported as means, or least squares means differences, with 95% confidence interval (95%CI).

#### 2 – ex vivo and in vivo electrophysiological data analyses

Friedman test followed by Dunn’s post-hoc test within groups were performed on electrophysiological data. Variations in cells properties were assessed using Kruskall-Wallis test or mixed ANOVA.

| Figure | | Number of rats | Comparison | Mean  differences | Confidence interval | Statistical test | p-Value |
| --- | --- | --- | --- | --- | --- | --- | --- |
| 1 | 1B | 8 EYFP long-access | Escalation (session 1 vs session 20) | 2.5 | 2 to 2.9 | post-hoc comparison for linear trend | F(1,123)=111, p<0.0001 |
|  | 1C | 8 EYFP long-access  7 EYFP short-access | Short vs Long-access:   - Baseline - Esc2 - Esc3 - Esc4     Long access :   - Esc4 vs Baseline - Esc4 vs Esc1 - Esc4 vs Esc 2 - Esc 4 vs Esc 3     Short access:   - Esc4 vs Baseline - Esc4 vs Esc1 - Esc4 vs Esc 2 - Esc 4 vs Esc 3 | 0.6  12.6  16.3  22.5      23.1  16.9  8.7  5      1.2  1.5  -1.3  -1.2 | -8.2 to 9.4  3.8 to 21.3  7.5 to 25  13.7 to 31.3      16.8 to 29.4  10.6 to 23.2  2.4 to 15  -1.3 to 11.3      -5.5 to 8  -5.2 to 8.3  -8 to 5.5  -8 to 5.6 | Sidak  Sidak  Sidak  Sidak      Dunnett  Dunnett  Dunnett  Dunnett      Dunnett  Dunnett  Dunnett  Dunnett | p>0.9999 p=0.0016 p<0.0001  p<0.0001      p<0.0001 p<0.0001 p=0.04  p=0.1588      p=0.9752 p=0.944 p=0.9714 p=0.9749 |
|  | 1D | 7 EYFP short-access | Without peer vs Baseline  Without peer vs Esc4  With peer vs Baseline  With peer vs Esc4  With peer vs Without peer | 1.8  0.9  -7.8  -8.7  -9.6 | -4.7 to 8.3  -5.7 to 7.5  -14.3 to -1.3  - 15.3 to -2  -16.1 to -3.1 | Tukey  Tukey  Tukey  Tukey  Tukey | p=0.8659 p=0.979 p=0.0153 p=0.008 p=0.0029 |
|  | 1E | 8 EYFP long-access | Without peer vs Esc4  Without peer vs Baseline  With peer vs Esc4  With peer vs Without peer  With peer vs Baseline | 0.3  23.2  -18.7  -19  4.3 | -9.6 to 10.8  13.4 to 33.1  -28.6 to -8.8  -28.9 to -9.1  -5.6 to 14.1 | Tukey  Tukey  Tukey  Tukey  Tukey | p=0.9998 p<0.0001 p=0.0002 p=0.0001 p=0.6882 |
|  | 1F | 8 EYFP long-access  7 EYFP short-access | Baseline short vs long access  Esc4 short vs long access  With peer short vs long access  Without peer short vs long access | -0,5964  -22,50  -12,56  -22,05 | -10,02 to 8,823  -31,92 to -13,08  -21,98 to -3,140  -31,47 to -12,63 | Sidak  Sidak  Sidak  Sidak | 0,9997  <0,0001  0,0046  <0,0001 |
| 2 | 2B | 8 EYFP long-access 9 ARCHT3.0 long-access | Escalation :  Esc4 vs Esc 1  Esc4 vs Esc2  Esc4 vs Esc3 | +35.9  +19.3  +12.7 | 23.3 to 48  7.1 to 31.4  0.5 to 24.8 | Dunnett  Dunnett  Dunnett | p<0.0001 p=0.0011 p=0.0401 |
|  | 2D | 7 EYFP short-access 6 ARCHT3.0 short-access | Baseline ARCHT3.0 vs EYFP  Esc4 ARCHT3.0 vs EYFP  Without peer ARCHT3.0 vs EYFP  With peer ARCHT3.0 vs EYFP | 3.8  0.3  0.1  8.6 | -12.1 to 4.5  -8.7 to 8  -8.2 to 8.4  0.2 to 16.9 | Sidak  Sidak  Sidak  Sidak | p=0.6725 p>0.9999 p>0.9999 p=0.0417 |
|  | 2E | 9 ARCHT3.0 long-access | Without peer vs Esc4  Without peer vs Baseline  With peer vs Without peer  With peer vs Baseline  With peer vs Esc4 | 18.6  -0.3  -3.8  -3.5  -22,4 | -31.2 to -6  -12.8 to 12.2  -16.3 to 8.6  -16 to 8.9  -34.9 to -9.9 | Tukey  Tukey  Tukey  Tukey  Tukey | p=0.0023 p=0.9999 p=0.8304 p=0.8604 p=0.0003 |
|  | 2F | 8 EYFP long-access 9 ARCHT3.0 long-access | Baseline ARCHT3.0 vs EYFP  Esc4 ARCHT3.0 vs EYFP  Without peer ARCHT3.0 vs EYFP  With peer ARCHT3.0 vs EYFP | -4.5  -0.3  17.8  3.2 | -17.6 to 8.5  13.4 to 12.7  -30.9 to -4.8-  9.9 to 16.2 | Sidak  Sidak  Sidak  Sidak | p=0.8491 p>0.9999 p=0.0035  p=0.963 |
| 3 | 2C | 9 hChR2 long-access  8 EYFP long-access  (no effect of the  optogenetic group) | Esc4 vs Esc1  Esc4 vs Esc2  Esc4 vs Esc3 | -40.7 -22  10 | -54.5 to -26.9 -35.8 to -8.2  -23.8 to -3.9 | Dunnett  Dunnett  Dunnett | p<0.0001 p=0.001 p=0.2061 |
|  | 2D | 9 hChR2 long-access | Without peer vs Esc4  Without peer vs Baseline  Without peer vs With peer  With peer vs Esc4  With peer vs Baseline | -19.53  -5.6  10.6  30.2  5.0 | 34.3 to -4.7  -20.4 to 9.1  -4.2 to 25.4  15.4 to 44.9  -9.8 to 19.8 | Tukey  Tukey  Tukey  Tukey  Tukey | p=0.0066 p=0.7227 p=0.2222 p<0.0001 p=0.786 |
|  | 2E | 9 hChR2 long-access  8 EYFP long-access | Baseline  Esc4  Without peer  With peer | -0.6  1.5  18.5  -9.7 | -13.7 to 12.5  -11.6 to 14.6  -31.6 to -5.4  -22.8 to 3.4 | Sidak  Sidak  Sidak  Sidak | p>0.9999 p=0.9972 p=0.0024 p=0.2239 |
| Supplemental | S2 | 9 ARCHT3.0 long-access  6 ARCHT3.0 short-access | Baseline  Esc4  Without peer  With peer | 8.9  23.2  4.1  0.8 | -5 to 22.8  9.2 to 37.1  -9.8 to 18  -13.1 to 14.8 | Sidak  Sidak  Sidak  Sidak | p=0.3567 p=0.0003 p=0.9092 p=0.9998 |

Supplementary Table 1: Statistical results of the behavioral paradigms

| Figure | | Number of cells | Parameters | Comparison | Rank sum  differences | Statistical test | p-Value |
| --- | --- | --- | --- | --- | --- | --- | --- |
| Supplemental | S4B | 11 ARCHT3.0 STN  transfected neurons *(ex vivo)* | 5 mW, pulse duration | 1s vs 10s  1s vs 15s  1s vs 20 | 32  43  51 | Dunn  Dunn  Dunn | p<0.01 p<0.0010 p<0.0010 |
|  | S4C | 11 ARCHT3.0 STN  transfected neurons *(ex vivo)* | 15s pulse at different light intensity | 2.5 mW vs 5 mW  2.5 mW vs 10 mW | 11  22 | Dunn  Dunn | p<0.05 p<0.001 |
|  | S4E | 10 hChR2 STN transfected neurons *(in vivo)* | 5 mW, pulse duration | 0.5s vs 4s  0.5s vs 5s | -42  -45 | Dunn  Dunn | p<0.001 p<0.001 |
|  | S4F | 10 hChR2 STN  transfected neurons *(in vivo)* | 1s pulse at different light intensity | 2.5 mW vs 10 mW  2.5 mW vs 20 mW | -19  -27 | Dunn  Dunn | p<0.01 p<0.001 |
|  | S5B | 10 ARCHT3.0 STN  transfected neurons *(ex vivo)* | STN neurons firing rate , light ON-OFF | 15s ON vs Baseline  30s OFF vs Baseline | 17  1 | Dunn  Dunn | p=0.0006 p>0.9999 |
|  | S5D | 7 ARCHT3.0 STN transfected neurons *(ex vivo)* | STN neurons firing rate , light ON-OFF | Laser bins 5 min ON vs Baseline Post-stimulation 5 min vs Baseline | -9  0 | Dunn  Dunn | p=0.0323 p>0.9999 |
|  | S5F | 16 ARCHT3.0 STN  transfected neurons *(in vivo)* | STN neurons firing rate , light ON-OFF | 15s ON vs Baseline  5s OFF vs Baseline | 26  4 | Dunn  Dunn | p<0.0001 p=0.959 |
|  | S5G | 16 ARCHT3.0 STN  transfected neurons *(in vivo)* | STN neurons firing rate , light ON-OFF | Laser bins 3 min ON vs Baseline Post-stimulation 1 min vs Baseline | 23  1 | Dunn  Dunn | p<0.0001 p>0.9999 |
|  | S6C | 10 hChR2 STN transfected neurons *(ex vivo)* | Member potential depending on light frequency stimulation | 10 Hz vs 100 Hz  10 Hz vs 130 Hz  10 Hz vs 150 Hz | 35  37  42 | Dunn  Dunn  Dunn | p<0.001 p<0.001 p<0.001 |
|  | S6E | 8 hChR2 STN transfected neurons *(ex vivo)* | STN neurons firing rate , light ON-OFF | Laser bins 5 min ON vs Baseline Post-stimulation 5 min vs Baseline | 9.5  1 | Dunn  Dunn  Dunn | p<0.0351 p>0.9999 |
|  | S6G | 16 hChR2 STN transfected neurons   - 7 non-responding cells - 9 responding cells *(in vivo)* | STN neurons firing rate , light ON-OFF | Non-responding cells:   - Laser bins 3 min ON vs Baseline - Post-stimulation 1 min vs Baseline     Responding cells:   - Laser bins 3 min ON vs Baseline - Post-stimulation 1 min vs Baseline | 5  4      -14  -1 | Dunn  Dunn      Dunn  Dunn | p=0.3629 p=0.5701      p=0.0019 p>0.9999 |

Supplementary Table 2: Statistical results of the electrophysiological recordings of STN neurons

**Supplementary results:**

REML results showed that hChR2 rats consumed a similar amount of cocaine than EYFP-control (optogenetic group effect: F_(1,15)_=0.01153, p=0.9159), and their consumption remained stable across the 5 sessions (session effect: F_(4,60)_=1.374, p=0.2539; optogenetic group x sessions: F_(4,60)_=2.27, p=0.0721). We then compared their cocaine consumption during the first 2h of the 20 long-access sessions with that of EYFP-control rats. REML results showed a significant effect of the sessions (F_(19,267)_=9.447, p<0.0001), but no effect of the optogenetic groups (F_(1,15)_=0.06603, p=0.8007), nor sessions x optogenetic groups (F_(19,267)_=1.001, p=0.4606). The number of cocaine injections taken in the six hours of each session was then averaged per block of 5 sessions (*‘Esc1’*, *‘Esc2’*, *‘Esc3’*, *‘Esc4’*) and compared between EYFP-control and hChR2 rats. REML results showed a significant effect of the 5-sessions blocks (F_(3,45)_=18.94, p<0.0001), but no effect of the optogenetic groups (F_(1,15)_=0.5088, p=0.4856), nor of their interaction (F_(3,45)_=0.104, p=0.9573). Dunnett’s post hoc analysis revealed that all animals progressively increased their cocaine intake (*‘Esc4’* vs. *‘Esc1’* mean difference: -40.7(95%CI: -54.5 to -26.9) injections, p<0.0001, vs. *‘Esc2’* mean difference: -22 (95%CI: -35.8 to -8.2), p=0.001, vs. *‘Esc3’* mean difference: -10 (95%CI: -23.8 to -3.9), p=0.2061).

We thus compared the drug intake across the five sessions of baseline, *‘Esc4’*, *‘Without peer’* and *‘With peer’* in hChR2 rats. REML results showed a significant effect of the conditions (F_(3,24)_=12.15, p<0.0001), but not of the sessions (F_(4,32)_=1.149, p=0.3517), nor of their interaction (F_(12,93)_=1.127, p=0.3486). Tukey’s post hoc analysis revealed that the laser stimulation was sufficient to decrease of 19.53 (95%CI:

-34.3 to -4.7) the number of cocaine injections compared to *‘Esc4’* (*‘Without peer’ vs. ‘Esc4’*, p=0.0066). Drug consumption reached a baseline level (*‘Without peer’ vs.* baseline mean difference: -5.6 (95%CI: -20.4 to 9.1), p=0.7227). The peer’s presence did not reduce the consumption further (*‘Without peer’ vs. ‘With peer’* mean difference: 10.6 (95%CI: -4.2 to 25.4) injections p=0.2222, *‘Esc4’ vs. ‘With peer’*: 30.2 (95%CI: 15.4 to 44.9), p<0.0001 and baseline *vs. ‘With peer’*: 5.0 (95%CI: -9.8 to 19.8), p=0.786).

Finally, we compared the mean number of cocaine injections taken during the baseline, Esc4 (the two first hours), *‘Without peer’* and *‘With peer’* was then compared between hChR2 and EYFP-control rats subjected to the long access procedure. REML results showed a significant effect of the optogenetic groups (F_(1,15)_=5.028, p=0.0405), of the conditions (F_(3,45)_=25.41, p<0.0001) and of their interaction (F(3,45)=9.764, p=0.0171). Sidak’s post hoc analysis revealed that hChR2 and EYF-control took a similar amount of cocaine injections when the laser was OFF (baseline mean difference: -0.6 (95%CI: -13.7 to 12.5) p>0.9999, *‘Esc4’* mean difference: 1.5 (95%CI: -11.6 to 14.6) p=0.9972). They also showed a similar consumption *‘With peer’* (mean difference: -9.7 (95%CI: -22.8 to 3.4) injections p=0.2239). However, *‘Without peer’*, hChR2 animals consumed 18.5 (95%CI: -31.6 to -5.4) cocaine injections than EYFP-control rats (p=0.0024).

**Supplementary figures :**

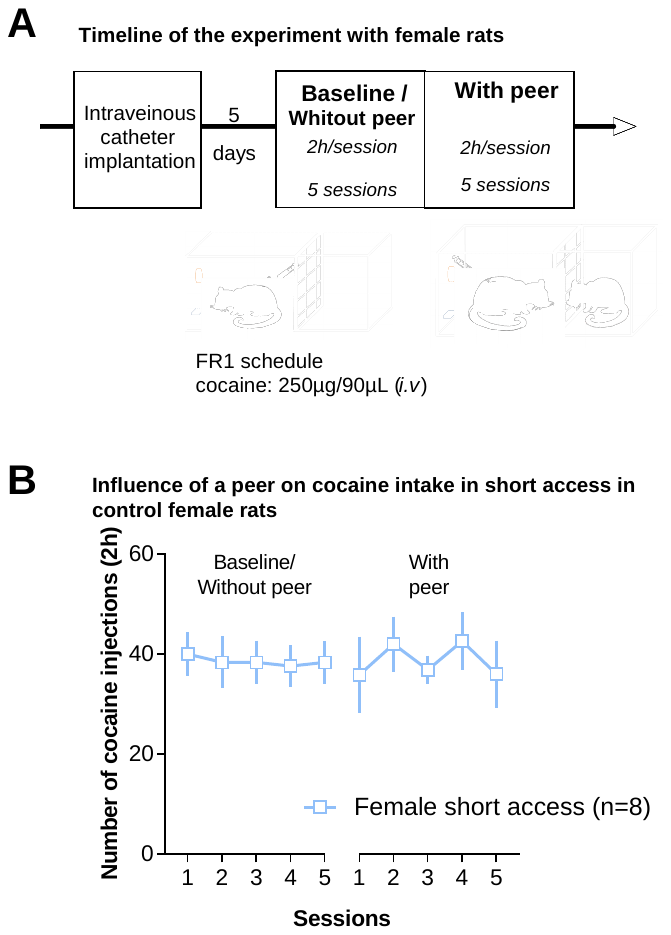

**Fig. S1: Female rats took a similar amount of cocaine across short access sessions, independently of the presence of a cocaine naive same-sex stranger peer.**

**(A)** Timeline of the experiment with female rats **(B)** Cocaine injections received by female rats maintained in the short access procedure (n=8) during the the baseline/ ‘*Without peer* sessions’ (38.55±0.39 injections) and ‘*With* peer’ (38.68±1.50) sessions

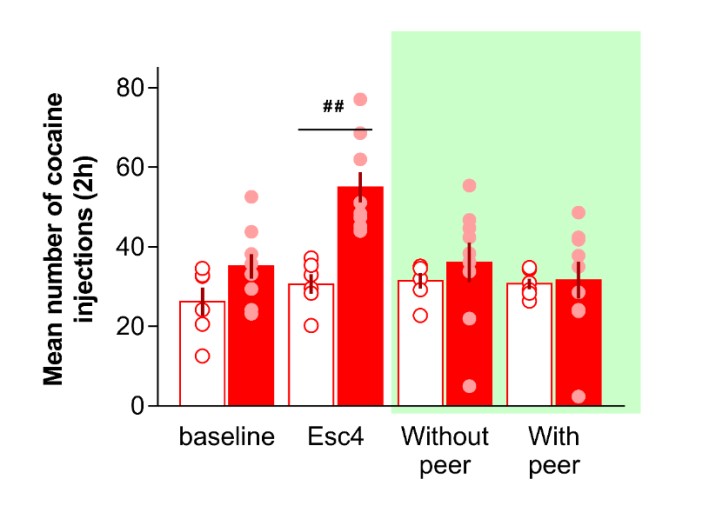

**Influence of a peer and STN inhibition on cocaine intake in short vs long access in ARCHT3.0 male rats**

**Fig. S2: All ARCHT3.0 rats took a similar amount of cocaine during the laser activation, independently of their drug escalation history.**

Mean number of cocaine injections received by ARCHT3.0 rats maintained in the short access procedure (n=6; empty red bars) and these subjected to the long access procedure (n=9; plain red bars) during the five 2h sessions of the baseline, Esc4 (the two first hours), ‘*Without peer*’ and ‘*With peer*’. Each dot represents an individual mean per block of 5 sessions. The laser activation is represented in green shaded area.

*## p<0.01 between short and long access.*

### Electrophysiological properties of STN neurons

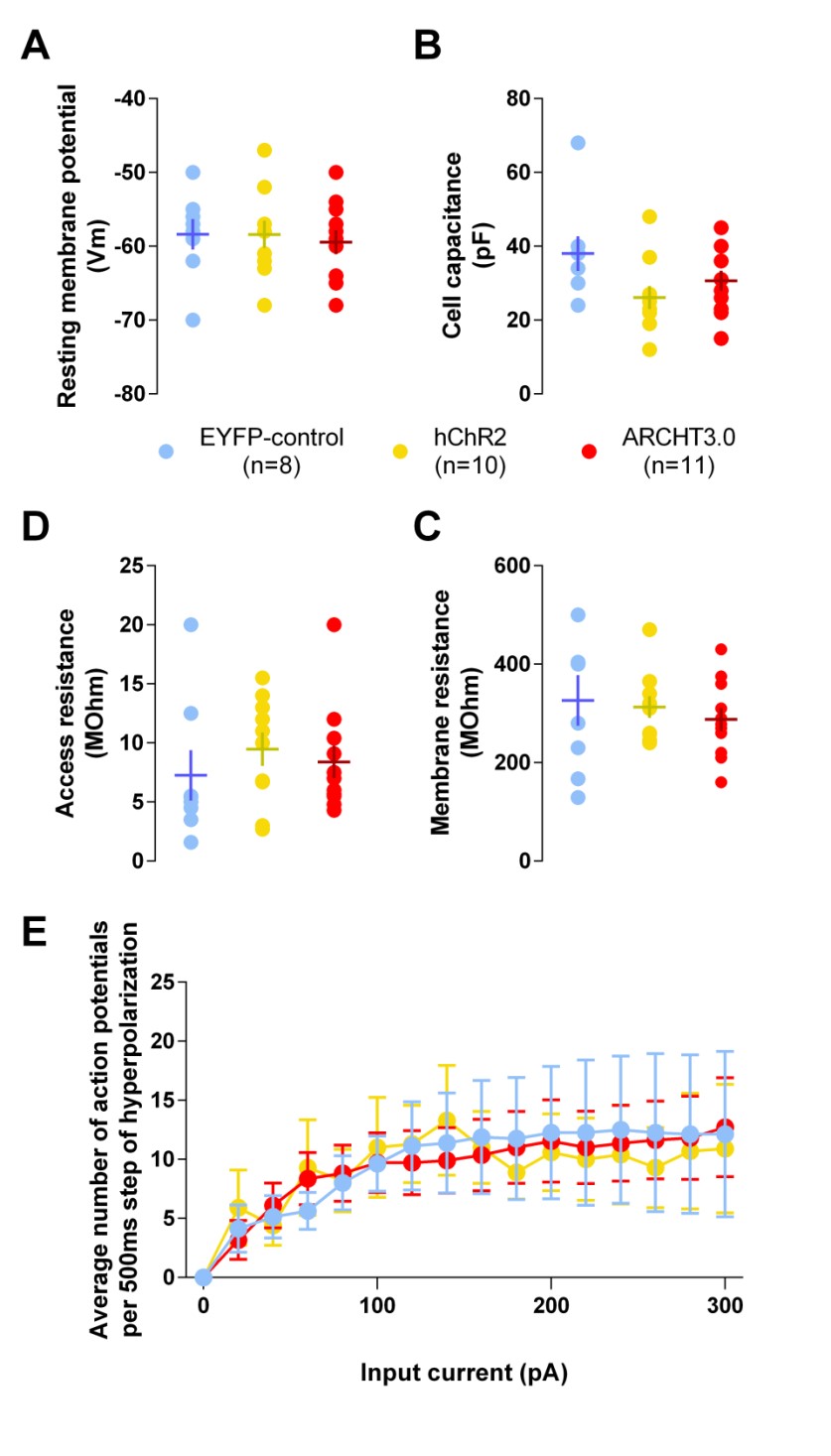

**Resting**

**P**

**otentiel**

**Cell Capacitance**

**Access Resistance**

**Membrane Resistance**

**Frequency Intensity curve**

**Fig. S3.** **Cells properties of STN neurons after viral infection recorded in whole cell configuration.**

Cell resting membrane potential (**A**), cell capacitance (**B**), membrane resistance (**C**), access resistance (**D**) and average number of action potentials (APs) following a 500ms step of hyperpolarization (**E**) for controls EYFP-control (n=8, blue), hChR2 (n=10, yellow) and ARCHT3.0 (n=11, red) cells.

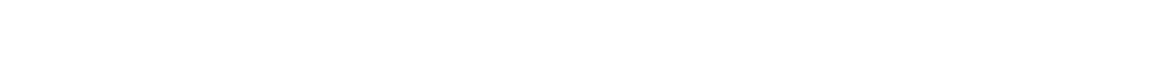

### Light induced currents in ARCHT STN infected cells

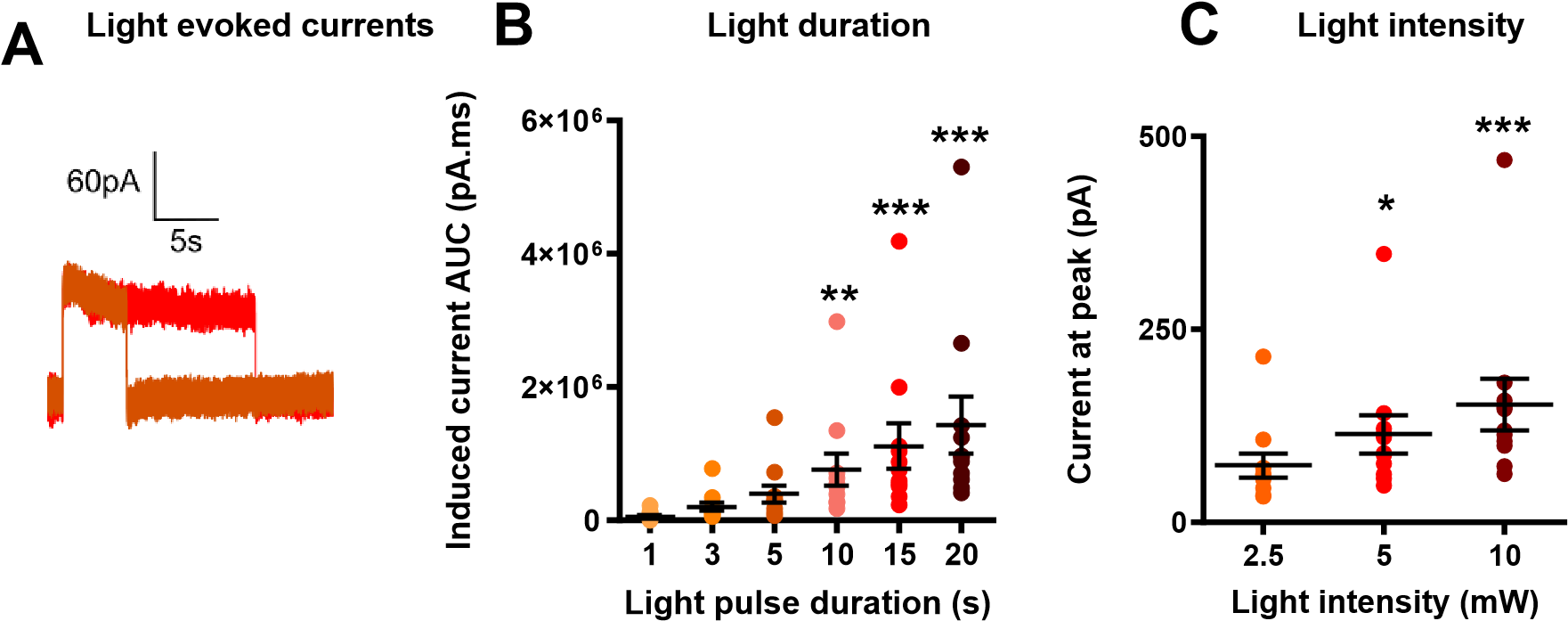

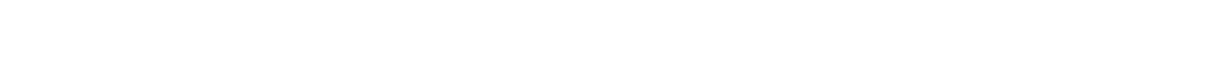

### Light induced currents in hChr2 STN infected cells

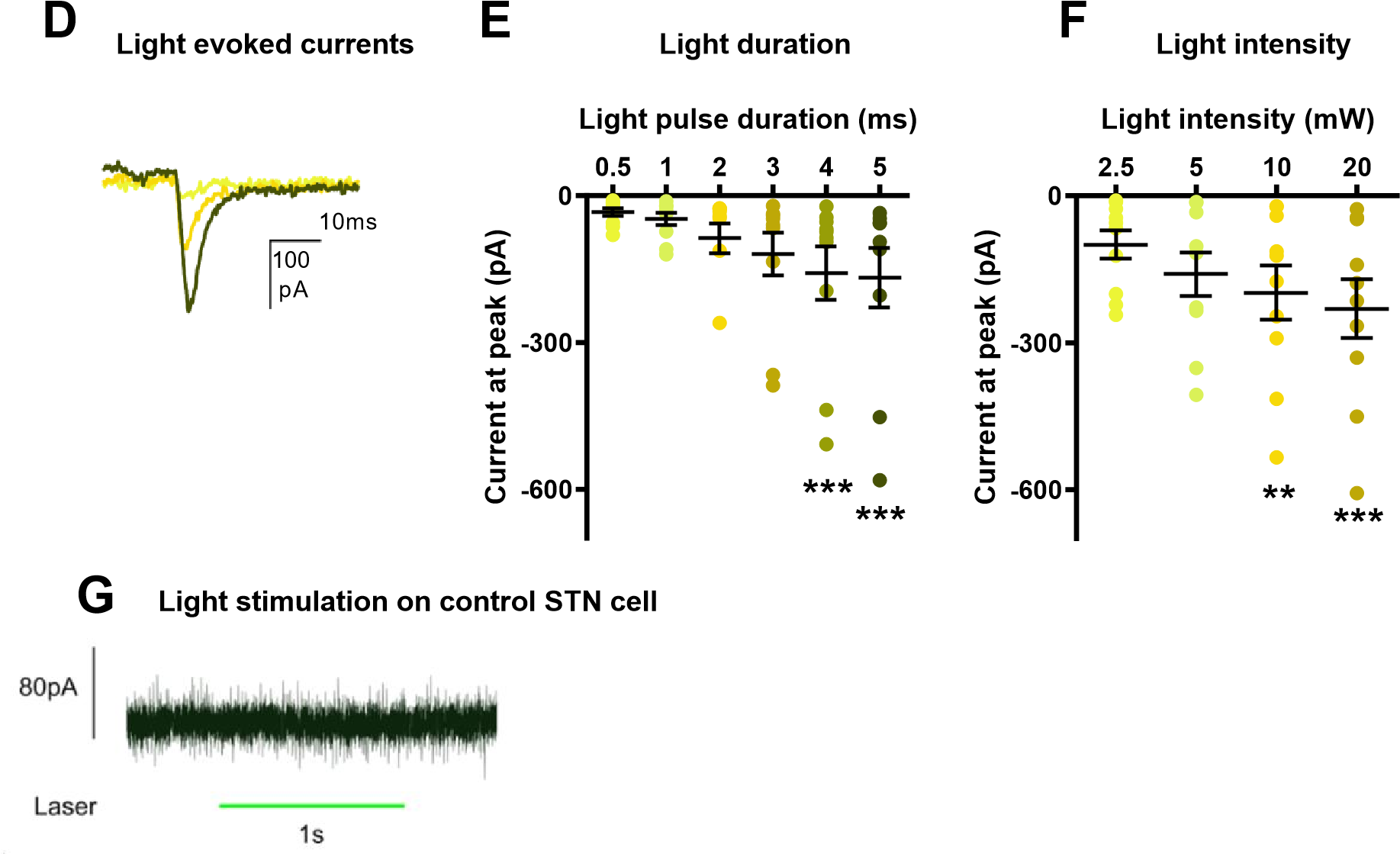

**Fig. S4. Evaluation of light stimulation parameters in STN neurons after viral infection.**

(**A**) Example of light induced inhibitory currents in one STN neuron expressing ARCHT3.0 by a 5s (brown) and 15s (blue) pulse. (**B**) Average area under the curve (AUC) calculated for different durations of light pulses with intensity set at 5mW in ARCHT3.0 cells (n=11). (**C**) Average current induced in ARCHT3.0 cells by a single 15s pulse applied with different light intensities. (**D**) Example of induced depolarizing current in hChR2 cells induced by a 0.5ms (pink), 2ms (brown) or 5ms (purple) light pulse. (**E**) Average induced current in hChR2 cells (n=10) depending on the duration of the light pulse with intensity set at 10mW. (**F**) Induced current in hChR2 cells by a 1s light pulse as a function of light intensity. (**G**) Typical response of a neuron expressing EYFP-control facing a 1s laser pulse at 20mW. ** p<0.05, ** p<0.01, ***p<0.001* compared with the lowest value of light parameter.

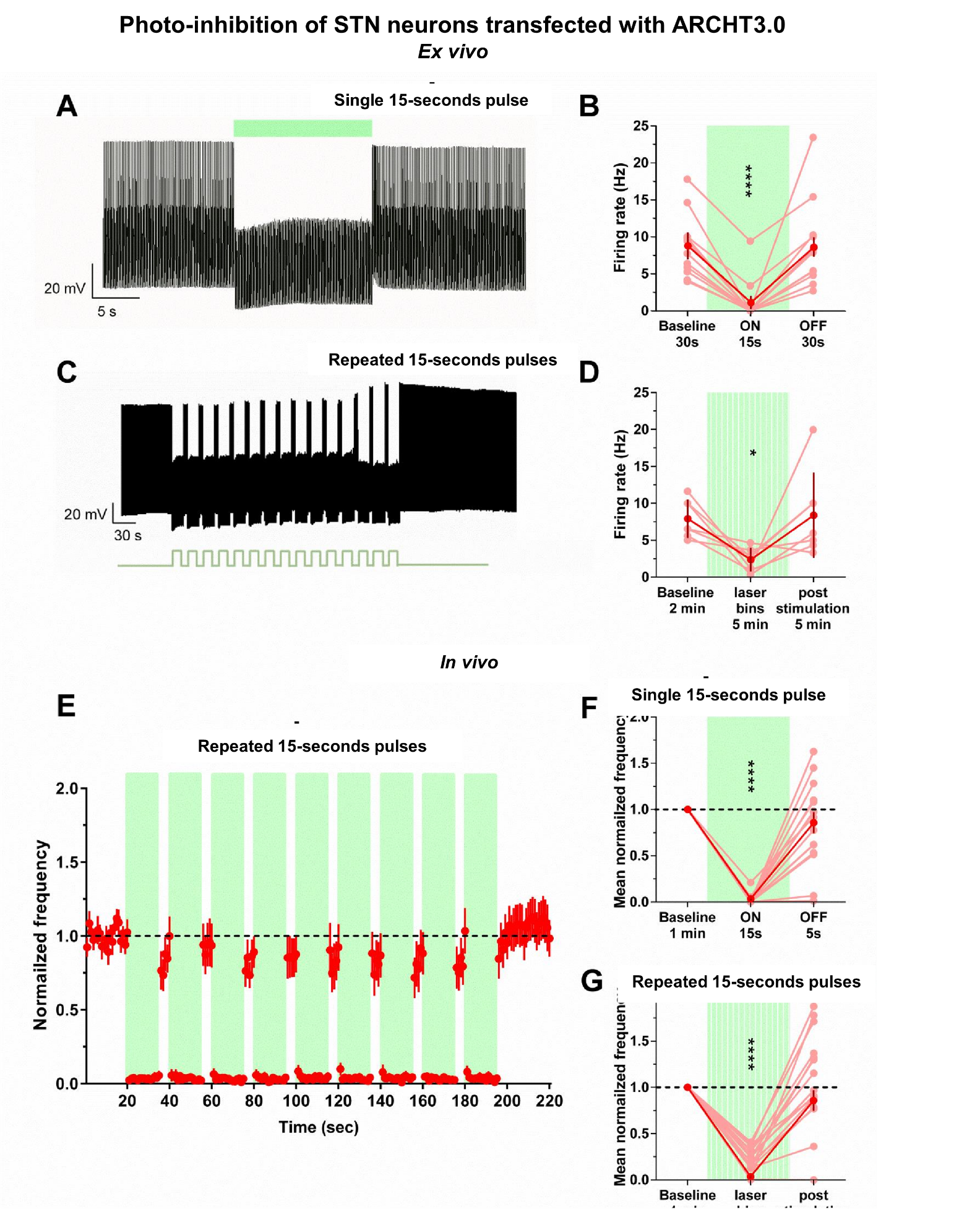

**Fig. S5 Electrophysiological assessment of STN optogenetic inhibition.**

**(A)** Typical response recorded in one STN neuron expressing ARCHT3.0 driven at a 10Hz frequency undergoing 15s of photo-inhibition (green line: laser ON). **(B)** Firing rate in ARCHT3.0-expressing STN neurons was reduced by a single 15s light pulse (n=11 cells). **(C)** Example of response for an ARCHT3.0-transfected STN neuron during 5 min of laser activation applied with the same pattern as during behavioral experiments (*i.e.* alternations of 15s ON, interleaved with 5s OFF, for 5min). **(D)** Firing rate in STN neurons expressing ARCHT3.0 was reduced by 5min of laser application at this pattern (n =7 cells). **(E)** Normalized frequency in ARCHT3.0-transfected STN neurons during 20-s of baseline recording, followed by 180-s of laser activation with the same pattern and 20s post stimulation (n=16 cells). **(F)** 15s laser activation (ON-15s), but not 5s of obscurity (OFF-5s) reduced the mean normalized frequency in ARCHT3.0transfected STN neurons, compared with their baseline activity (n=16 cells). **(G)** 180-s of laser bins diminished the mean normalized frequency in STN neurons, compared with their baseline activity. Following 180-s of laser bins, these cells returned to their baseline level of activity. Full green area corresponds to continuous laser activation. Hatched green area corresponds to laser bins. In **B,D,F,G**: Each light red dot and line represents an individual value. Dark red dot and lines correspond to the mean and SEM.

**: p<0.05,****:p<0.0001.*

### Photo-inhibition of STN neurons transfected with hChR2

***Ex vivo***

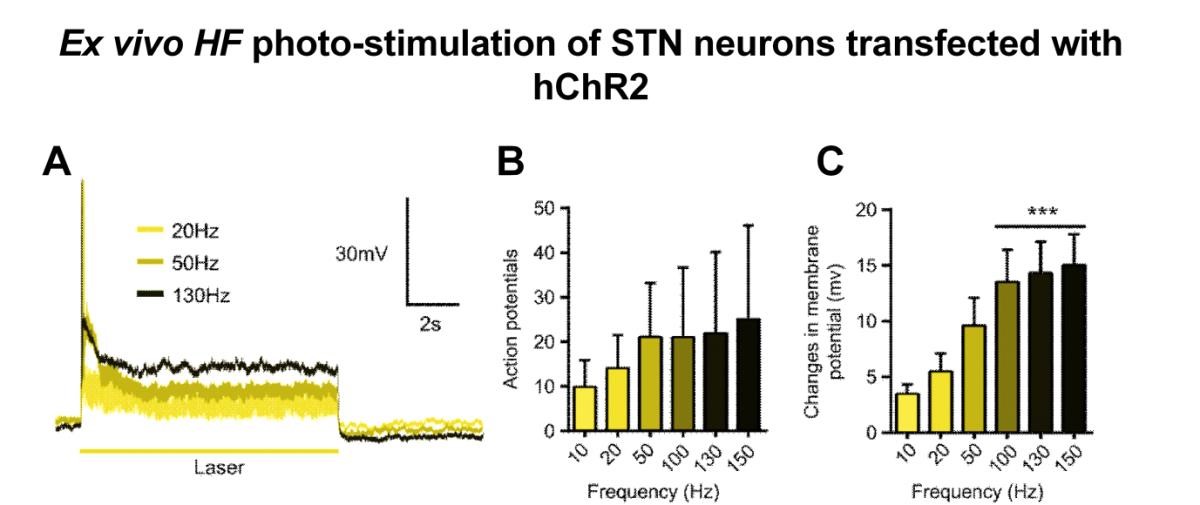

**Light evoked currents**

**Frequency influence**

**on Action Potential**

**Membrane**

**potential changes**

**130 Hz 5-minutes light stimulation**

***In vivo***

**130 Hz 3-minutes light stimulation**

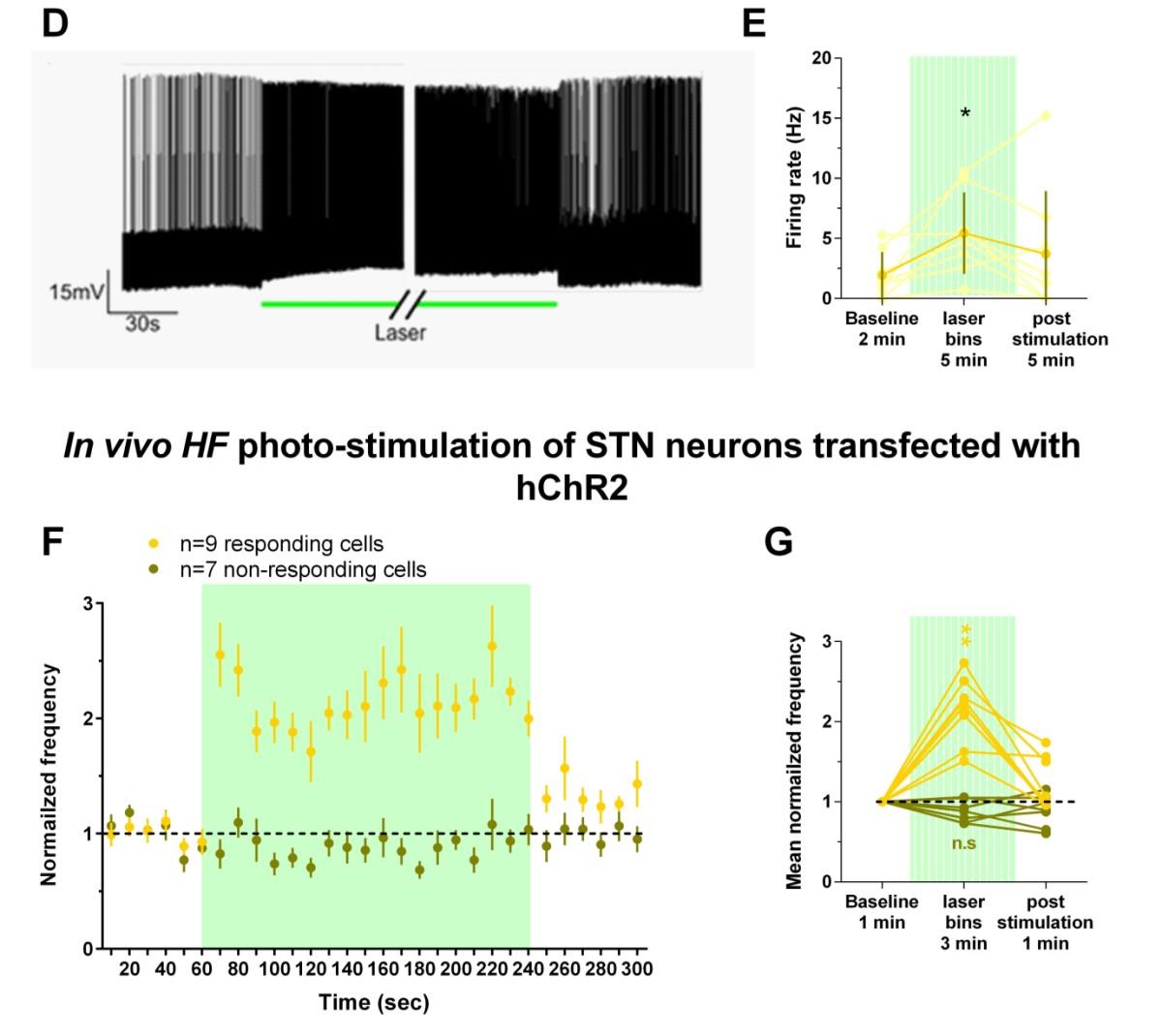

**Fig. S6 Electrophysiological assessment of STN optogenetic HF-stimulation**.

(**A**) Typical excitatory response recorded in current-clamp for a STN neuron expressing hChR2 and facing 10s of laser stimulation at various frequencies: 20 (light yellow), 50 (dark yellow) and 130Hz (black), eliciting a few action potentials but increasing membrane potential. (**B**) Number of action potentials elicited during 10s of photo-stimulation (n=10 cells). (**C**) Variation of membrane potential caused by the 10s optogenetic stimulation at various frequencies (n=10 cells). (**D**) Example of response in hChR2 positive STN neuron at onset (left panel) and offset (right panel) of 5min laser stimulation at 130Hz (green line) used during behavioral testing. (**E**): Average changes in firing rate induced by 5 min of 130Hz laser stimulation (n=8 cells). Light dot and lines represent individual values. Dark line and dot correspond to the mean and SEM. (**F**) Normalized frequency in responding (light yellow; n=9) and nonresponding (dark yellow; n=7) hChR2-expressing STN neurons during 1min of baseline recording, followed by 180s of HF photo-stimulation (at 130Hz) and 1 min post stimulation. (**G**) Mean normalized frequency during baseline, HF photo-stimulation and post-stimulation in responding and non-responding cells.

Full green area corresponds to 130Hz laser stimulation.

**: p<0.05, **:p<0.01, ***:p<0.001.*
